# The complex evolution of the metazoan HSP70 gene family

**DOI:** 10.1101/2020.09.21.307264

**Authors:** Er-meng Yu, Tatsuki Yoshinaga, Frank L. Jalufka, Hashimul Ehsan, David B. Mark Welch, Gen Kaneko

## Abstract

The metazoan 70-kDa heat shock protein (HSP70) family contains several members localized in different subcellular compartments. The cytosolic members have been classified into inducible HSP70s and constitutive heat shock cognates (HSC70s), but their distinction and evolutionary relationship remain unclear because of occasional reports of “constitutive HSP70s” and the lack of cross-phylum comparisons. Here we provide novel insights into the evolution of these important molecular chaperones. Phylogenetic analyses of ∼100 full-length HSP70s revealed an ancient duplication that gave rise to two lineages from which all metazoan cytosolic HSP70s descend. One lineage (A) contains a relatively small number of Lophotrochozoan and Ecdysozoan genes, none of which have been shown to be constitutively expressed (i.e., either inducible or unknown). The other lineage (B) included both inducible and constitutive genes from diverse phyla. Species-specific duplications are present in both lineages, and Lineage B contains well-supported phylum-specific clades for Rotifera, Nematoda, and Chordata. Some genes in Lineage B have likely independently acquired inducibility, which may explain the sporadic distribution of “HSP70” or “HSC70” in previous analyses. Consistent with the diversification history within each group, inducible members show lower purifying selection pressure compared to constitutive members. These results illustrate the evolutionary history of the HSP70 family, encouraging us to propose a new nomenclature: “HSP70 + subcellular localization + linage + copy number in the organism + inducible or constitutive, if known.” e.g., HSP70cA1i for cytosolic Lineage A, copy 1, inducible.

## Introduction

The 70-kDa heat shock protein (HSP70) family members play important roles in various cellular processes including heat shock response, folding of newly synthesized proteins, protein transport, and protein degradation. These apparently diverse functions are attributed to their chaperone activity (Daugaard et al. 2007; Hartl et al. 2011), by which they prevent the aggregation and misfolding of target proteins. Typically, HSP70 family members bind to denaturing or newly synthesized proteins by recognizing up to ten hydrophobic amino acid residues exposed to the protein surface because of the misfolding. The release of the HSP70 members, which is triggered by ATP hydrolysis, facilitates the proper folding of the target proteins (Zuiderweg et al. 2017). The HSP70 family contains organelle-specific members localized in cytosol, endoplasmic reticulum (ER), mitochondria, and chloroplasts (Miernyk 1997), and these organelle-specific types not only perform chaperoning functions in the organelles but are also known to contribute to protein transport across organelle membranes.

HSP70 family members are often upregulated upon various stresses that disrupt protein folding, such as heat treatment, exposure to toxic materials, ultraviolet irradiation, and pathogen attack (Sørensen et al. 2003; Baird et al. 2006). The upregulation is primarily regulated at the transcription level by transcription factors called heat shock factors (HSFs), particularly HSF1. Under unstressed conditions, HSF1 remains in a monomeric state. Various stresses lead to the formation of HSF1 trimers, which in turn bind to heat shock elements (HSEs) in the promoter region of HSP70 family member genes and promote their transcription. The HSF1-dependent transactivation system is generally conserved among eukaryotes, although species-specific differences in HSF1 function have been reported in many organisms such as the fruit fly *Drosophila melanogaster* (Jedlicka et al. 1997; Marchler and Wu 2001) and yeast *Saccharomyces cerevisiae* (Sorger and Pelham 1988). The expression of HSP70 family member is also regulated by many other mechanisms such as the unfolded protein response, chromatin modification, and other transcription factors depending on the type of stress and subcellular localization (De Nadal et al. 2011; Garbuz 2017). HSP70 family member genes have therefore been used as biomarkers of environmental stresses both in laboratory and field experiments (Sanders 1993; Ceyhun et al. 2010; Judge et al. 2011).

On the other hand, several cytosolic HSP70 family members show constitutive expression patterns. These proteins are involved in the folding of newly synthesized proteins and are traditionally called 70-kDa heat shock cognates (HSC70). However, the distinction between HSP70s and HSC70s remains unclear because of the occasional reports of “constitutive HSP70s,” especially from invertebrates (Jayasena et al. 1999; Piano et al. 2002; Liu et al. 2017). Stress-induced upregulation of HSC70 has also been reported in several animals including shrimps (Luan et al. 2010), snails (Zheng et al. 2012), and fish (Yabu et al. 2011). Furthermore, little is known for the evolutionary relationship between metazoan HSP70s and HSC70s because of the lack of cross-phylum comparisons. A previous phylogenetic analysis indicated that metazoan HSP70 family members can be classified into invertebrate HSP70s, vertebrate HSP70s, and HSC70s from both vertebrates and invertebrates (Kourtidis et al. 2006), but this study comprises only a few phyla. In-depth phylogenetic analysis on HSP70 family members with a broad sampling would offer further insight into the classification of this important group of molecular chaperones.

In this study, we sought to trace the evolutionary history of metazoan HSP70 family members by cross-phylum phylogenetic analyses with a particular attention to their stress inducibility. The specific hypothesis was that the cross-phylum analysis would provide the essential information to solve the evolutionary relationship between metazoan HSP70s and HSC70s. In order to add insights from an emerging model in evolutionary biology to the analyses, we first cloned two stress-inducible HSP70 family member genes from a monogonont rotifer, *Brachionus plicatilis*. The phylum Rotifera, composed of mostly microscopic aquatic animals with about 1,000 cells and a ciliated head structure, is part of the Gnathifera, a group of basal-branching phyla related to Lophotrochozoa, which contains Mollusca and Annelida. Together these groups form a sister-clade to Ecdysozoa, which contains the established invertebrate models *Caenorhabditis elegans* and *D*. *melanogaster* (Struck et al. 2014; Fröbius and Funch 2017). Genetic information from Gnathifera remains limited, making rotifers useful models in evolutionary studies. Our results added novel groups of stress-inducible HSP70, providing important insight into the evolutionary history of the metazoan HSP70 family.

## Results

Molecular Characterization of Rotifer HSP70 Genes A combination of 3’ and 5’ RACE identified two *B. plicatilis* HSP70 genes consisting of 2,121 and 2,248 bp, designated as HSP70cB1i and HSP70cB2i, respectively (see Discussion for the nomenclature). The nucleotide sequence of the open reading frames, as well as the absence of introns, were ascertained by single PCR experiments using cDNA or genomic DNA, respectively, followed by DNA sequencing. The deduced amino acid sequences of HSP70-1 and HSP70-2 genes were more than 99% identical to each other (supplementary fig. S1). Both contain the HSP70 protein family signatures, IDLGTTYS, IFDLGGGTFDVSIL, and IVLVGGSTRIPKVQK (Prosite motifs PS00297, PS00329, and PS01036, respectively), as well as the non-organelle stress protein motif RARFEEL found in several cytosolic HSP70s (Lo et al. 2004; Cottin et al. 2008; Simoncelli et al. 2010; Zheng et al. 2012). The major difference between *B. plicatilis* HSP70 genes was the number of GGMP repeats, which are involved in binding to the HSP70-HSP90-organizing protein (Hop), a cofactor of HSP70 (Demand et al. 1998). The nucleotide sequences of *B. plicatilis* HSP70cB1i and HSP70cB2i genes were registered into the DDBJ/EMBL/GenBank databases with accession numbers AB775784 and AB775785, respectively. The two HSP70 genes were also found in the *B. plicatilis* genomic sequence.

We next examined whether the mRNA levels of *B. plicatilis* HSP70s are increased by heat stress. Due to the high sequence identity between *B. plicatilis* HSP70 genes and difficulty in designing a TaqMan probe, we employed semi-quantitative RT-PCR to measure their respective mRNA levels. Primers were designed to amplify DNA fragments of 112 and 124 bp from HSP70cB1i and HSP70cB2i cDNAs, respectively (fig. 1A), and the PCR products were separated using a polyacrylamide gel. The mRNA levels of HSP70cB1i and HSP70cB2i in heat-treated rotifers were 2.9 and 7.5 times higher, respectively, than those in control rotifers (fig. 1B and 1C). These results indicate that both genes encode heat-inducible HSP70.

**FIG. 1.**
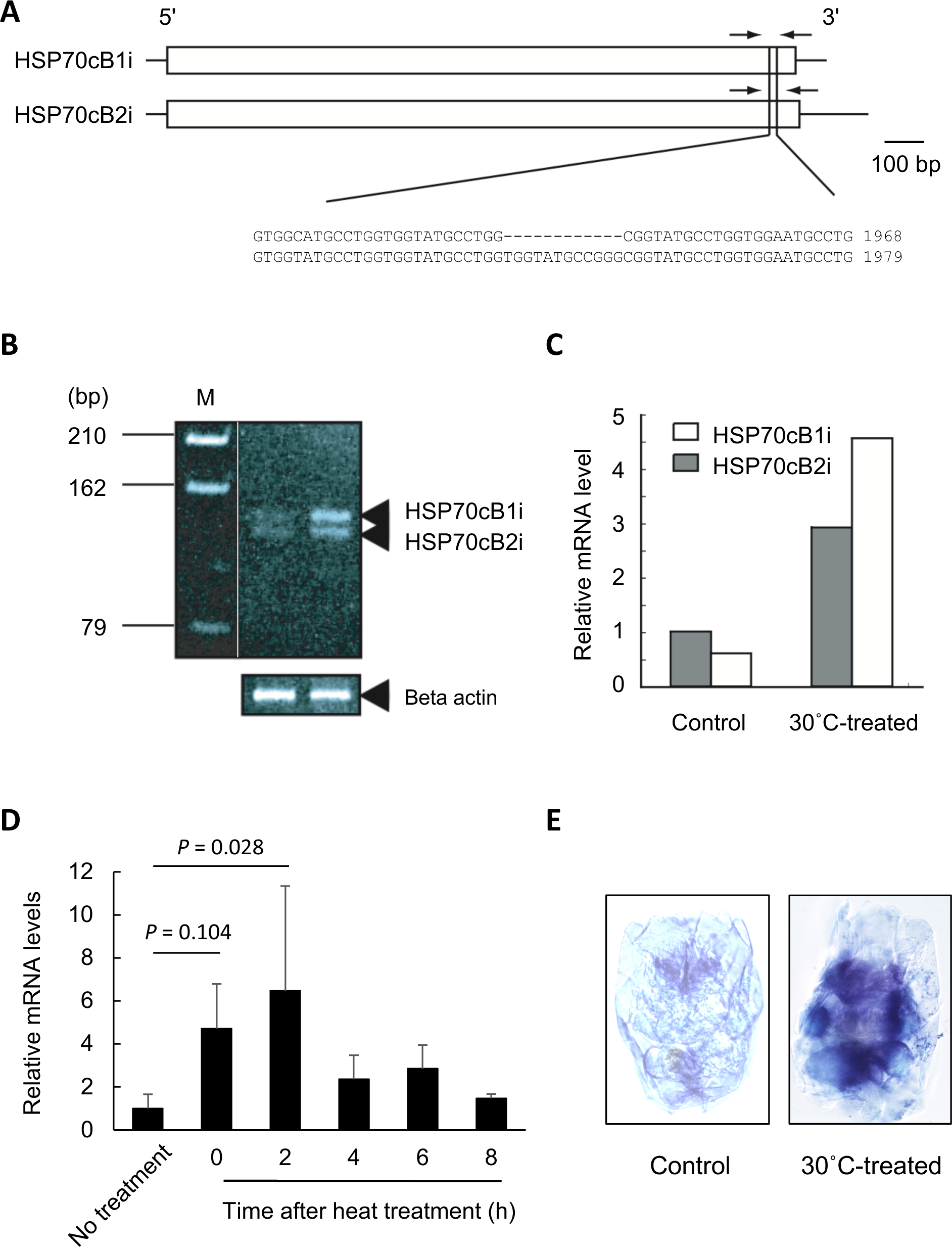
Molecular characterization of *Brachionus plicatilis* HSP70 genes. (A-C) Semi-quantitative reverse transcription-PCR. A representative result is shown out of three independent experiments. Location of primers rHSP70_gapF and rHSP70_gapR that amplify cDNA fragment of 112 and 124 bp from HSP70cB1i and HSP70cB2i cDNAs, respectively (A). Polyacrylamide gel electrophoresis patterns of the RT-PCR product. Cycle numbers within a linear range of PCR amplification were determined to be 24 to 28 cycles for both cDNAs by preliminary experiments on the basis of signal intensities of amplified products by RT-PCR (B). Signal intensities of HSP70cB1i and HSP70cB2i genes standardized to those of b-actin (C). (D) Heat stress-induced expression of HSP70 quantified by real-time PCR. Beta-actin gene was used as the internal control. Bars represent standard errors from three replications. One-way ANOVA detected significant effects of heat treatment (F = 3.281, df = 5, *P* = 0.0426). Dunnett’s multiple comparison was used to detect significant differences between the control and other groups. Difference between control and two hours group was statistically significant (t = 3.246, *P* = 0.028). (E) *In situ* hybridization for *B. plicatilis* HSP70 genes. Rotifers were fixed 4 h after the heat treatment.

Since the two *B. plicatilis* HSP70 genes showed a similar expression pattern in response to heat stress, we used quantitative real-time PCR primers that amplify both HSP70cB1i and HSP70cB2i genes in the subsequent time-course expression analyses (fig. 1D). The relative mRNA levels were increased by heat stress at 40°;C for 10 min and reached at the maximum level 2 h after the heat stress. The difference in mRNA levels between the control and 2 h groups was statistically significant. The mRNA levels of *B. plicatilis* HSP70s remained twofold 8 h after the heat stress. We then performed *in situ* hybridization using DIG-labeled RNA probes (fig. 1E). The probe region contained only five base pair differences between the two HSP70 genes, and thus the RNA probes were likely to hybridize with both HSP70cB1i and HSP70cB2i mRNAs. The hybridization signals increased in all tissues of heat-treated rotifers. Altogether, these results indicate that the *B. plicatilis* HSP70cB1i and HSP70cB2i genes are heat-inducible and have similar expression profiles.

Phylogenetic Analysis of HSP70 Family Members

To assess the relationship of the *B. plicatilis* HSP70 genes with those of other rotifers and other metazoans, we constructed gene trees of more than 100 HSP70 family members, including members associated with the mitochondria and ER and many sequences known only from automated genome annotation as “HSP70.” To assess the robustness of the results we used alignments made by three different methods [Clustal-Omega (Sievers et al. 2011), M-Coffee (Wallace et al. 2006), and Expresso (Armougom et al. 2006)] to construct consensus trees with maximum-likelihood (RAxML) and Bayesian (MrBayes) approaches. The RAxML PROTGAMMAAUTO function found that the LG model of protein evolution (Le and Gascuel 2008) was optimal for all three alignments and this model was used to construct ML and Bayesian consensus trees. The six trees were in general agreement for major nodes; fig. 2 shows the Bayesian consensus tree of the Expresso alignment and support for major nodes from each approach is shown in table 1. In all analyses HSP70 family members known to be mitochondria- and ER-specific formed two monophyletic groups (nodes 11 and 12 in fig. 2) in agreement with previous reports (Boorstein et al. 1994; Miernyk 1997; Nikolaidis and Nei 2004). All family members known to be cytosolic formed a third group (node 1; clades 1–6 in fig. 2), with yeast cytosolic HSP70s (clade 6) basal to all metazoan cytosolic HSP70s. The tree topology generally supported the species phylogeny. Within the metazoan cytosolic HSP70s (node 2; clades 1–5) there were two lineages: Lineage A was well supported and contained of genes from arthropods, molluscs, and rotifers (node 3; clade 1); Lineage B was more poorly resolved and represented by many diverse vertebrate and invertebrate phyla (node 4; clades 2–5). There are two groups of vertebrate cytosolic HSP70s (nodes 8 and 9), which form a monophyletic group (node 7; clade 5) in Bayesian and RAxML analyses of Expresso and Clustal alignments. Unexpectedly, we found two additional clusters of cytosolic HSP70 family members in all analyses: one composed of HSP70 genes from the phylum Rotifera (node 5; clade 2) and the other from the phylum Nematoda (node 6; clade 3). This suggests the evolution of two unique subgroups of HSP70, distinct from conventional vertebrate and invertebrate HSP70s. We also constructed phylogenetic trees mainly using HSP70 family members of known expression patterns (supplementary figs. S2 and S3). Both of the two HSP70 lineages as well as most of the nodes described above were identified, supporting the robustness of the tree shown in fig. 2, although several clades were poorly resolved in supplementary figs. 2 and 3 that have limited number of samples.

**FIG. 2.**
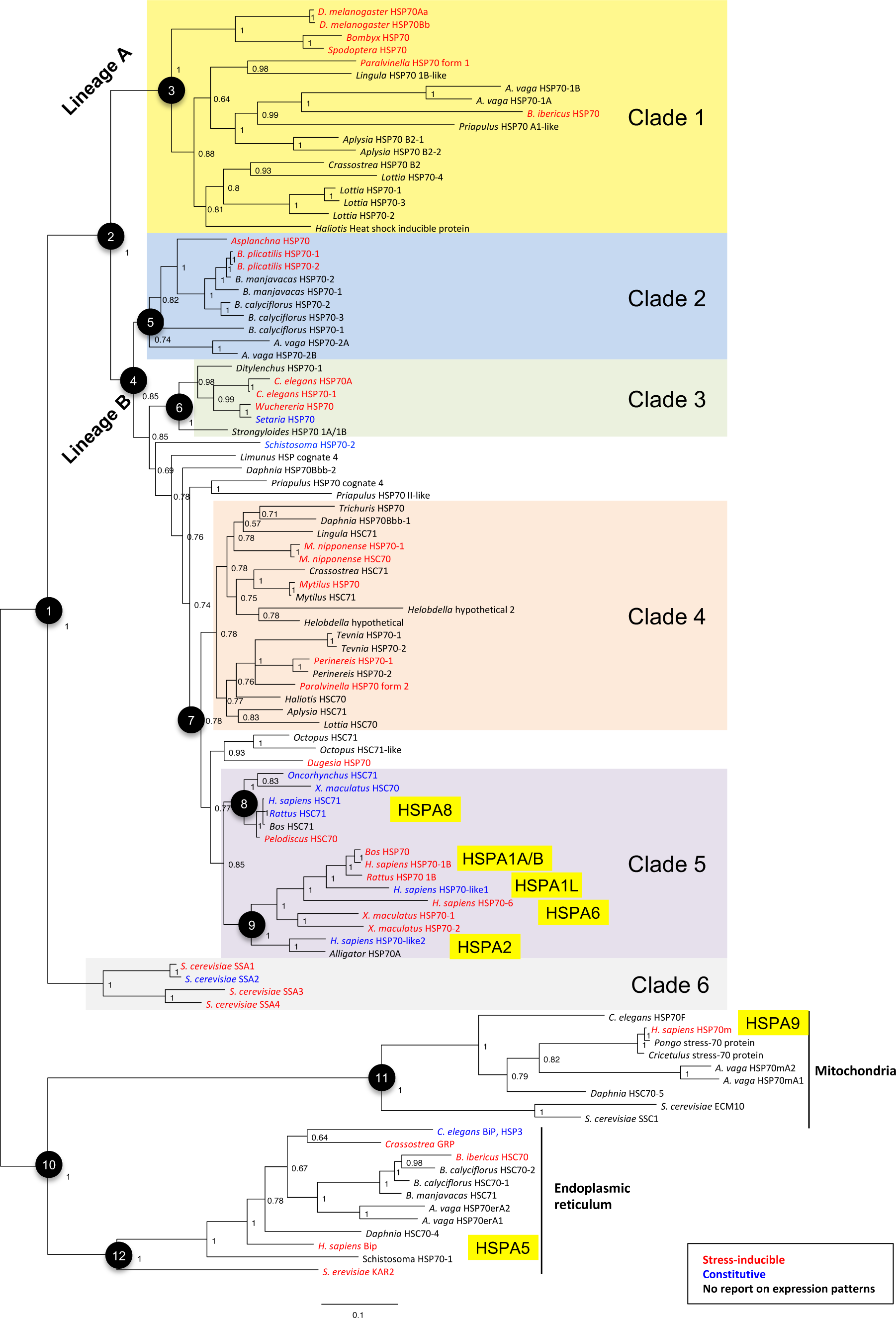
Bayesian consensus tree of HSP70 family members. Nodes discussed in the text are indicated. Numbers on the branches indicate the posterior probability support for each node. Stress- inducible and constitutive genes are shown in red and blue, respectively. No data on the stress inducibility is available for genes shown in black. Approved HGNC names of human HSP70 family members are shown in yellow boxes for reference. The DDBJ/EMBL/GenBank accession numbers and other information are summarized in supplementary table S2. Clades 1 to 6 include only cytosolic members of the HSP70 family.

**FIG. 3.**
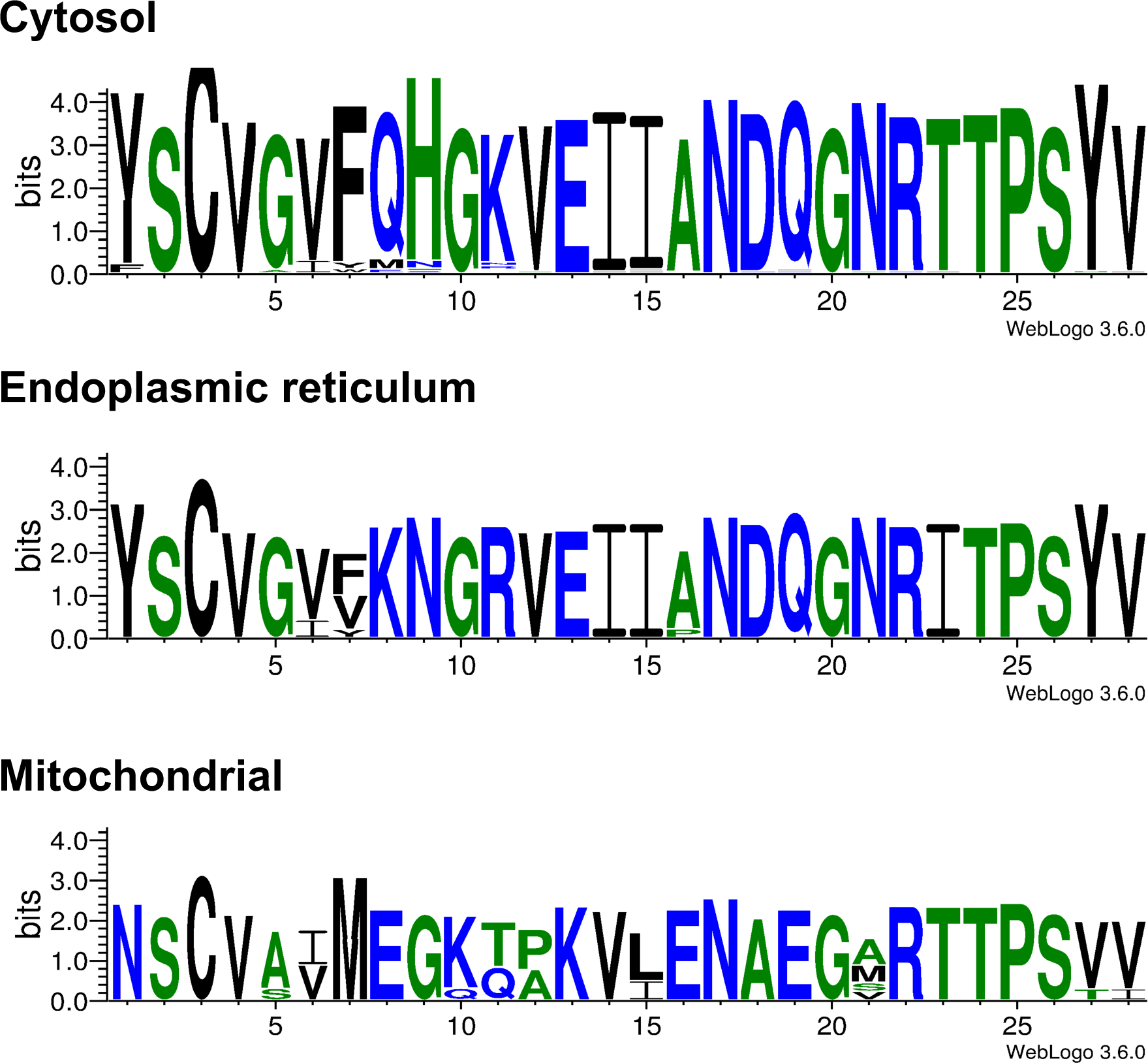
Logo visualization of mitochondrial-, endoplasmic reticulum-, and cytosolic-specific sequence near the N terminus of the HSP70 family members. Compared to the cytosolic form, ER- specific forms have several conserved amino acid substitutions: Q8K, H9N, K11R, and T23I. Similarly, mitochondrial forms have Y1N, G5A, F7M, Q8E, H9G, G10K, K11T/Q, V12P/A, E13K, I14V, I15L, A16E, D17A, and Q18E.

**Table 1.** Support for major nodes from each approach

|  |  | ClustalOmega |  | Expresso |  | M-Coffee |  |
| --- | --- | --- | --- | --- | --- | --- | --- |
|  |  | MrBayes | RAxML | MrBayes | RAxML | MrBayes | RAxML |
| 1 | Cytosolic | 100 | 99 | 100 | 99 | 100 | 100 |
| 2 | Metazoan | 100 | 95 | 100 | 99 | 100 | 94 |
| 3 | Lineage 1 | (a) | 99 | 100 | 99 | 100 | 100 |
| 4 | Lineage 2 | (b) | 41 | 85 | 67 | (b) | (c) |
| 5 | Rotifer | 100 | 48 | 74 | 48 | 100 | 65 |
| 6 | Nematode | 94 | 93 (d,e) | 100 (e) | 96 (e) | 100 (e) | 86 (e) |
| 7 | Chordate |  | 42 | 85 | 35 |  |  |
| 8 | HSC | 100 | 91 | 100 | 86 | 100 | 31 |
| 9 | HSP | 100 | 88 | 100 | 83 | 100 | 87 |
| 10 | Outgroup | 100 | 99 | 100 | 99 | 100 | 100 |
| 11 | Mitochondrial | 100 | 100 | 100 | 97 | 100 | 100 |
| 12 | ER | 100 | 99 | 100 | 100 | 100 | 100 |

(a) Rotifer genes (node 5) is included in node 3 with 83% support
(b) Rotifer genes (node 5) sister group to node 3 with 80% support, other paraphyly
(c) Nematode genes (node 6) basal; rotifer genes sister group to node 3 with 46% support, other paraphyly
(d) 36% with Trichuris (immediate outgroup to node 6)
(e) Trichuris not in node 6

The phylogenetic breadth of our analyses provided the opportunity to search for sequence motifs characteristic of different groups within the HSP70 family. We identified a single region near the N terminus (but after signal peptide motifs) that discriminates between metazoan mitochondrial, ER, and cytosolic HSP70s (fig. 3), which enables better discrimination for organelle-specific HSP70s together with the well-known C-terminal motifs for cytosolic (EEVD) and ER (HDEL/KDEL) HSP70s (supplementary fig. S4). We also found the RARFEEL motif, previously described as possibly cytosolic-specific in conjunction with the EEVD motif (Cottin et al. 2008; Simoncelli et al. 2010; Zheng et al. 2012), in several invertebrate genes that are found with high confidence in the ER-associated node 12 (fig. 2; *B. ibericus* HSC70, *C. gigas* GRP, and *B. manjavacas* HSC71). This result suggests that the RARFEEL is not a reliable predictor of cytosol association. Indeed, this motif was identified by a comparison of few sequences (Lo et al. 2004) and appears to have no strong scientific support.

Stress-inducible invertebrate HSP70s are reported to have an extra serine residue in their ATPase domain (Kourtidis et al. 2006; Garbuz et al. 2011). In our analysis this insertion occurs only in Lineage A irrespective of the stress-inducibility, where it is present in all sequences except *B. ibericus* HSP70 and one of four *Lottia* HSP70s; in *Priapulus* HSP70-1 the serine has been replaced by alanine (fig. 4). These results suggest that the extra serine residue is a characteristic of this lineage and may be associated with the stress-inducibility of the ancestral HSP70, which will be tested by broader sampling focusing on this residue. The alignment result supports the idea that HSP70s from Nematoda and Rotifera evolved separately from the those in Annelida, Mollusca and Arthropoda, which have been generally recognized as “invertebrate HSP70s”.

Synonymous and Nonsynonymous Substitution Rates

Lastly, we calculated the synonymous and nonsynonymous substitution rates of 34 representative cytosolic HSP70 member genes. We primarily selected genes with known expression patterns (stress-inducible or constitutive) from each clade, hypothesizing that constitutive HSP70 family members have been under stronger purifying selection compared to stress-inducible members. Only well-conserved regions were used for the calculation (supplementary fig. S5). We noted that *B. ibericus* HSP70 (GU574486) has an insertion at nucleotide position 403 and a nucleotide deletion at position 563, causing a frame shift. Deduced amino acid sequences encoded by nucleotides 403–563 were markedly different from those of other HSP70 family members because of the frame shift, which resulted in the long genetic distance in phylogenetic trees (fig. 2). Because the frame shift greatly affects the synonymous and nonsynonymous substitution rates, this sequence was excluded from our analysis.

For the selected 34 sequences, we calculated synonymous and nonsynonymous substitution rates (Ks and Ka) for all combinations of representative cytosolic members of the HSP70 family from metazoans (fig. 5A). Synonymous substitution rates (Ks) were saturated in many combinations and only a limited number of combinations produced meaningful results (229 out of 465 combinations, supplementary material 1). The Ka/Ks values calculated for the 229 combinations were averaged for each clade (fig. 5B). Overall, Ka/Ks values were markedly lower than 1, indicating that cytosolic HSP70 family member genes have been under purifying selection. The purifying selection pressure appears to have been less in Lineage A (clade 1) than in Lineage B (clades 2-5). The effect size between these lineages ranged from 0.19 to 1.64 (fig. 6C). Since the effect size larger than 0.8 indicate large difference in multiple comparisons, these results support our hypothesis that constitutive HSP70 family members [clade 4 and clade 5 (node 8)] have been under stronger purifying selection.

**FIG. 5.**
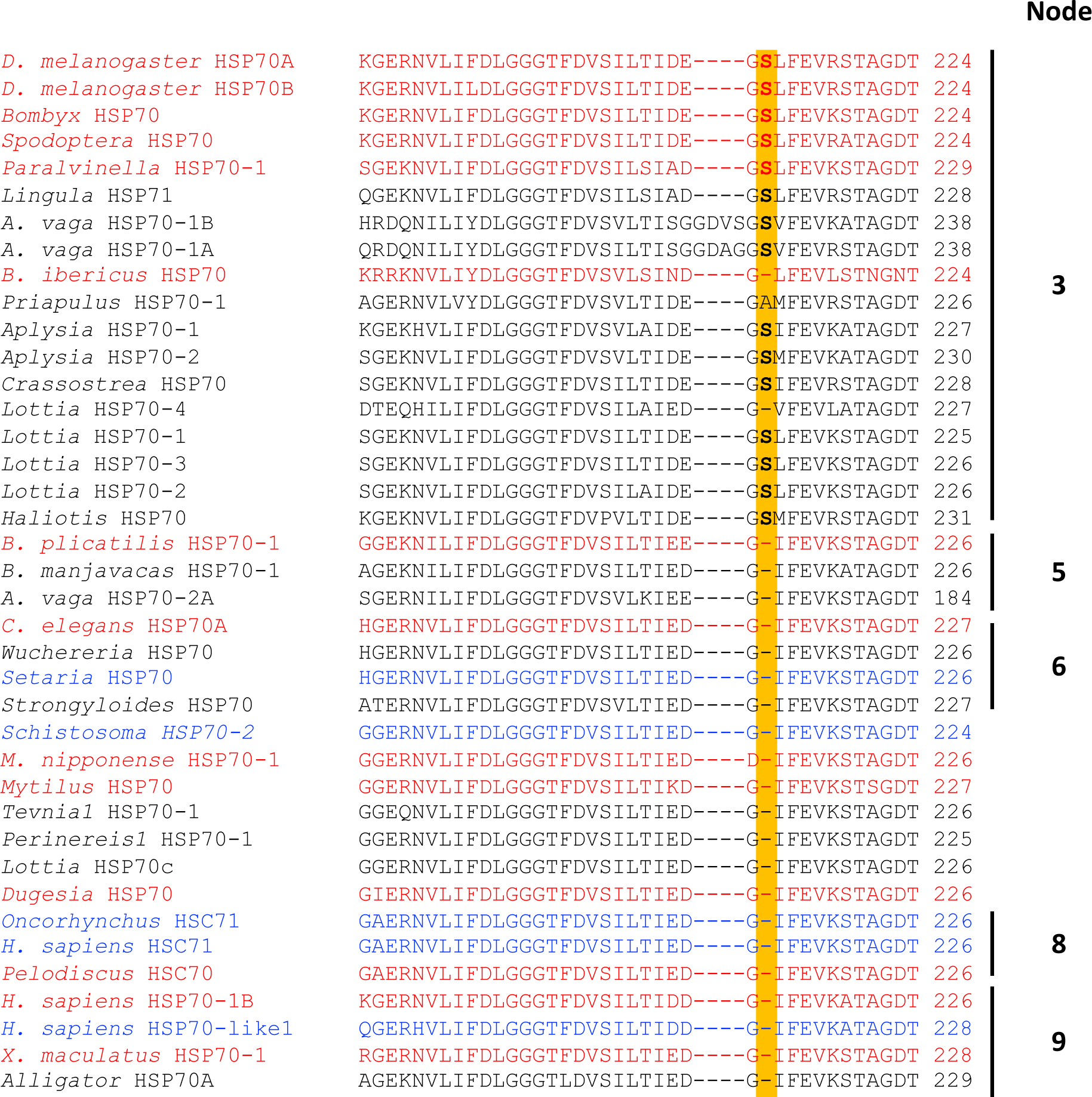
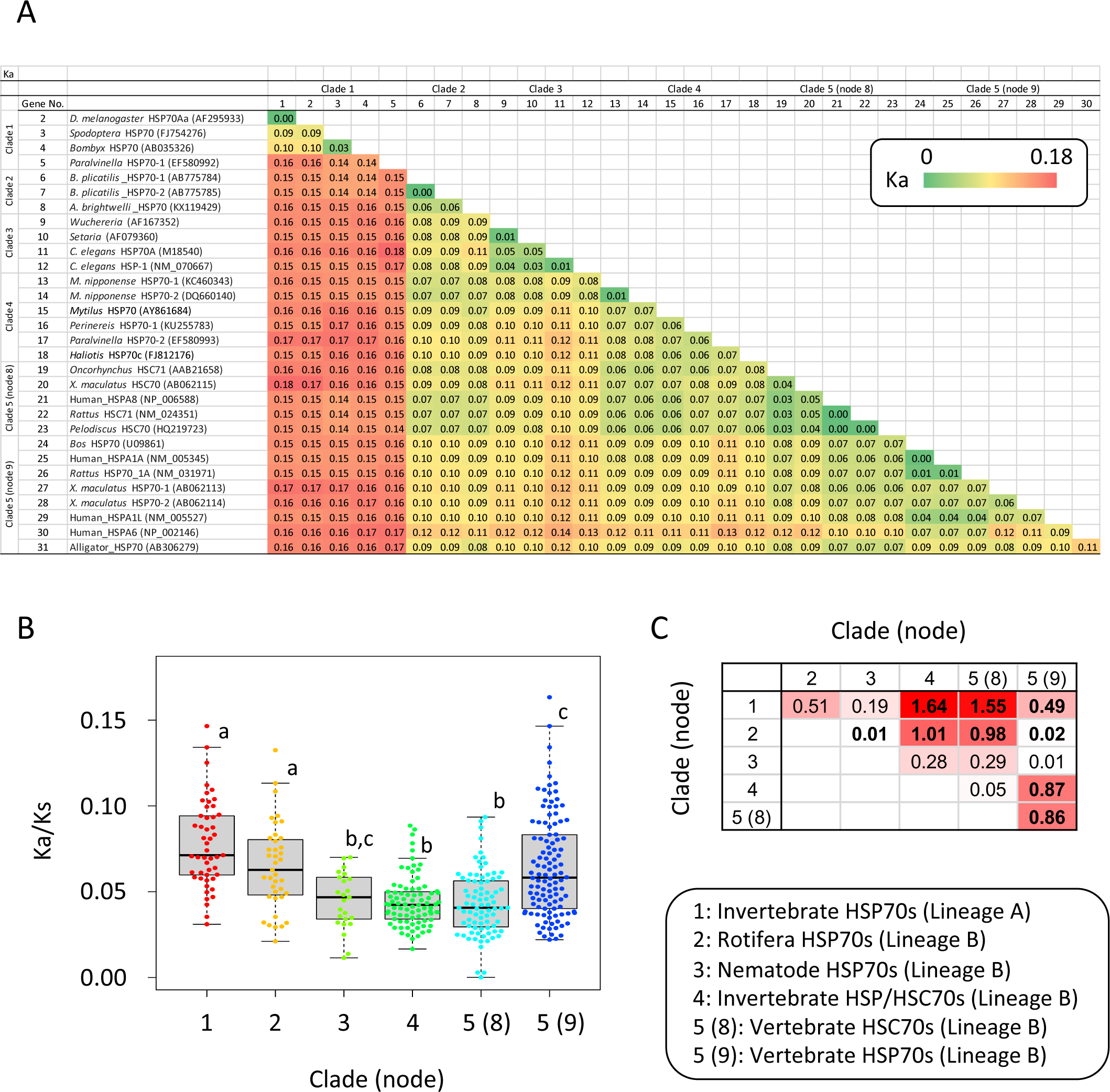
Comparison of the amino acid sequences of the HSP70 family members from invertebrates. Nodes in the phylogenetic tree (fig. 3) are indicated on the right margin, and serine residues specific to node 3 are highlighted. Stress-inducible and non-inducible genes are shown in red and blue, respectively. No data on the stress inducibility is available for genes shown in black. The accession numbers and other information are summarized in supplementary table S2.

**FIG. 6.**
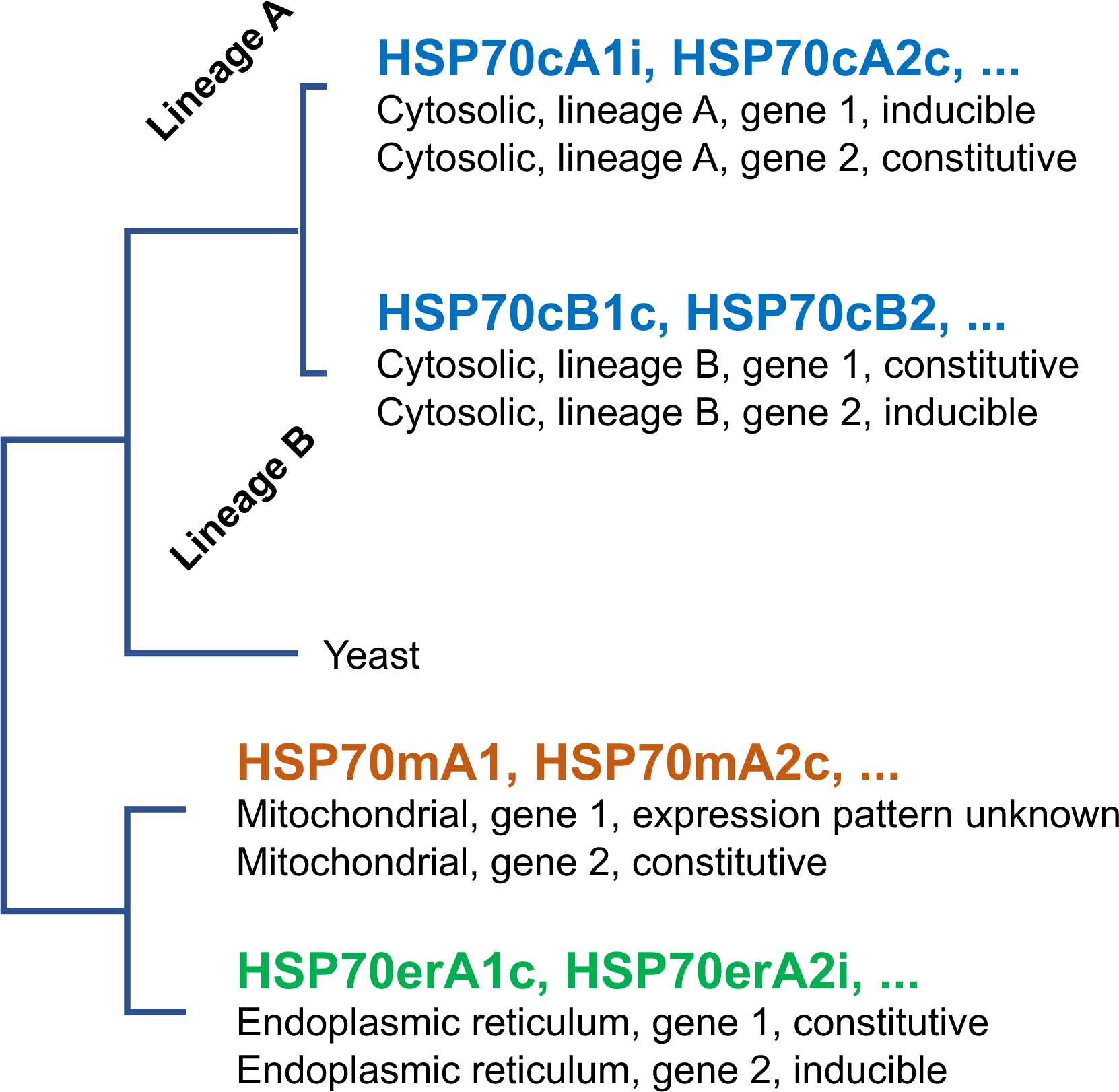
Selection pressure on cytosolic HSP70 family member genes. (A) Nonsynonymous substitution rates (Ka values) between 34 HSP70 family member genes. Gene number 1 is the fruit fly HSP70 Bb (AF295957) gene. The substitution rate of fruit fly HSP70 Aa and Bb gene was 00395526, which is shown as 0.00 in the figure. Accession numbers for other genes are indicated in the figure. Clades and nodes correspond to those in fig. 2. (B) Bee swarm boxplots of Ka/Ks values for each clade. Only Ka/Ks values calculated from inter-cluster pairs were averaged. Statistical differences were calculated by Kruskal-Wallis test (chi-squared = 88.704, df = 5, p < 2.2e-16) followed by the non-parametric post-hoc tests (pairwise Wilcox test with P value adjustment by the Holm method). Clades sharing same letters are not significantly different at the 5% level of significance. One outlier in clade 3 (0.53) is not included in the plot (see supplementary material 1) although this value was used for all statistical analyses. (C) Effect sizes (Cohen’s d) for all comparisons. Combinations where significant differences were found in the pairwise Wilcox test are shown in bold.

Within Lineage B, the selection pressure was strongest in Nematode HSP70s (clade 3), invertebrate HSC70/HSP70s (clade 4), and vertebrate HSC70s (clade 5, node 8). In particular, vertebrate HSC70s (clade 5, node 8) and HSP70s (clade 5, node 9) showed significant difference in the Ka/Ks with a large effect size of 0.86 despite their close phylogenetic relationship. These results also support the above hypothesis. To confirm how much this finding depends on the sampling, we conducted the same analysis with a larger sample size, including most genes used in the phylogenetic analysis, and obtained consistent results (supplementary fig. S6).

## Discussion

A previous phylogenetic analysis by Kourtidis et al. (2006) classified metazoan cytosolic HSP70s into three groups, invertebrate HSP70, vertebrate HSP70, and HSC70 including both vertebrate and invertebrate orthologues using data from a few taxa (Kourtidis et al. 2006). The monophyletic origin of HSC70s was also proposed by Nikolaidis and Nei (2004) based on their analysis using *Drosophila* and nematode sequences (Nikolaidis and Nei 2004). Surprisingly, to our knowledge, there has been no significant update on the evolution of metazoan HSP70 family after these small-scale studies despite the continuing expansion of available genomic data. Here we show that there are at least two types of ancestral cytosolic HSP70 family member genes in all analyzed phyla of metazoans, one (node 3; clade 1) giving birth to Lineage A of invertebrate HSP70s, and a second (node 4; clade 5) to Lineage B of both vertebrate and invertebrate HSP70s and all HSC70 genes. This second lineage has further diversified within diverse phyla: Rotifera (node 5; clade 2), Nematoda (node 6; clade 3), and Chordata (node 7; clade 5) (fig. 2), which has not been identified in the previous reports. The tree topology is generally consistent with the species phylogeny, indicating that the effect of sequence contamination is very limited.

Inter-clade comparison of the synonymous and nonsynonymous substitution rates shows that while all HSP70 family members are under strong purifying selection, the pressure to conserve amino acid sequence varies across clades. The presence of purifying selection has been reported for mammalian (Hess et al. 2018), nematode (Nikolaidis and Nei 2004), and molluscan (Kourtidis et al. 2006) HSP70 genes, and our cross-phylum analysis was consistent with these reports. Most dramatically, the canonical HSC70 genes of vertebrates (node 8; clade 5) showed stronger purifying selection pressure than the HSP70 family members in node 9 (clade 5). This concept can be further generalized by thoroughly investigating the saturation rate between each HSP70 gene (Pollock and Larkin 2004). It is thus speculated that the role of HSC70 protein, the folding of newly synthesized proteins, is more essential for cell survival compared to the role of the HSP70 cognates (node 9; clade 5). In line with this speculation, cells deficient in the HSC70 gene are non-viable (Florin et al. 2004), whereas HSP70 knockout mice are mostly viable and fertile, although they are known to be sensitive to stress (Daugaard et al. 2007). Meanwhile, genetic distance between each clade is not uniform, and the gene conversion may also affect the accuracy of this calculation. In this regard our data should be interpreted in a semiquantitative manner.

It is known that nematode, mammalian, and molluscan HSP70 families have experienced gene conversion events (Nikolaidis and Nei 2004; Kourtidis et al. 2006; Hess et al. 2018). In addition to the purifying selection, gene conversion also likely contributes to the highly conserved nature of the HSP70 family members found in this study. The tree topology might have been affected by the gene conversion in closely related paralogues and orthologues such as human HSPA1A and HSPA1B (Hess et al. 2018). However, gene conversion generally functions to conserve the sequence in a species, and thus the overall topology of the metazoan tree is unlikely to be significantly affected. Gene conversion rate should be about 10 times higher than the mutation rate to significantly affect the topological distance according to a genomic simulation study (Touchon et al. 2009), but the nucleotide identity between sequences is often lower than 70% in our phylogenetic analysis.

Given the presence of inducible and constitutively expressed genes in many clades of metazoan cytosolic HSP70, it is difficult to predict the ancestral state of this phenotype, and whether a gene is inducible or constitutive expression does not predict its relationship to other family members. Phylogenetic analyses, the novel signature sequences near the N terminus (fig. 4), and the extra serine residue (fig. 5) would provide better clues for the classification of HSP70 family members than their expression patterns under stress.

Based on the largest-ever cross-phylum phylogenetic analysis on HSP70 family, we propose a new nomenclature that reflects the phylogenetic relationship of the HSP70 family members (fig. 6). The proposed name of an HSP70 family member protein consists of “HSP70 + subcellular localization + linage + copy number found in the organism + inducible or constitutive, if known.” For example, HSP70cA1i represents a cytosolic HSP70 in Lineage A, copy 1, inducible; and HSP70mA1 for mitochondrial linage of unknown inducibility. Although the current study found only one lineage in mitochondrial and ER members, the use of “A” in these organelle-specific isoforms would be beneficial. Systematic lineage names such as HSP70cD and HSP70mB can be assigned for novel lineages that may be discovered in future. Gene names are represented by lowercase italics of protein names (e.g., *hsp70cb2i* and *hsp70era1*).

In conclusion, the present study identified two novel HSP70 family members from a monogonont rotifer, an emerging model in evolutionary biology. The subsequent phylogenetic analyses illustrated the evolutionary history of the metazoan HSP70 family, in which stress inducibility does not reflect evolutionary history. The proposed nomenclature based on molecular evolution will help us understand the true nature of the HSP70 family.

## Materials and Methods

### Culture

We used the *B. plicatilis* Ishikawa strain (also called *Brachionus* sp. ISKW), originally isolated from a Japanese eel culture pond (Yoshinaga et al. 2004), for cDNA cloning, semi-quantitative RT-PCR, quantitative real-time PCR, and *in situ* hybridization. The cytochrome oxidase subunit I gene sequence has been registered for detailed identification of this strain (GenBank accession number LC422762). Rotifers were cultured at 25 °;C by a standard protocol (Kaneko et al. 2016) using half-diluted Brujewicz artificial seawater and fed the algae *Chlorella regularis* (Nikkai Center, Tokyo, Japan) at a final concentration of approximately 7 × 10^6^ cells/mL. See supplementary material 2 for details of molecular characterization of rotifer HSP70 genes.

### Phylogenetic Analysis

Metazoan HSP70 family genes were selected based on the following criteria: 1) genes from diverse metazoan phyla are included; 2) HSP70 genes with known expression patterns are prioritized; 3) both Lineage A and Lineage B genes from the same organism are included when possible; and 4) the number of genes from a single species is limited to ∼5 because within-phylum gene duplications do not change the tree topology. Annotated and unannotated genome and transcriptome databases were used. A fasta file containing sequences used for the tree is provided as supplementary material 3.

A multiple sequence alignment of HSP70 and HSC70 amino acid sequences was produced using Clustal Omega, Expresso, and M-Coffee with default parameters. Bayesian phylogenetic analysis was performed using MrBayes (v3.2.6) on the CIPRES scientific gateway v3.3 (Miller et al. 2010) with four chains each for 10^7^ generations. Every 1000th trees were sampled, and the first 25% of samples were discarded as a burn-in. Bayesian consensus trees were visualized with the TreeView software v1.4.3.

### Synonymous and Nonsynonymous Substitution Rates

R version 4.0.4 was used for the calculation of the synonymous and nonsynonymous substitution rates (the kaks function), Kruskal-Wallis test (kruskal.test function), and post-hoc test (pairwise.wilcox.test function) on the Macintosh platform. Heat maps were created in Microsoft Excel 2016 (Redmond, MA).

## Author Contributions

Conceived and designed the experiments: TY, GK. Performed experiments and data analyses: EMY, FLJ, DMW, GK. Contributed reagents/materials/analysis tools: EH, DMW, GK. Wrote the paper: DMW, GK. Read and approved the final manuscript: EMY, TY, FLJ, EH, DMW, GK.

## Acknowledgements

We express our sincere thanks to Dr. Kristin Gribble, Marine Biological Laboratory, for helpful discussions and comments on the manuscript. The authors are grateful to Cynthia Rodriguez and Jamie Wilson, University of Houston-Victoria, for their help in gene expression analyses. GK was supported by funding from M.G. and Lillie A. Johnson Foundation, Victoria, Texas.

---

**Table 1.**
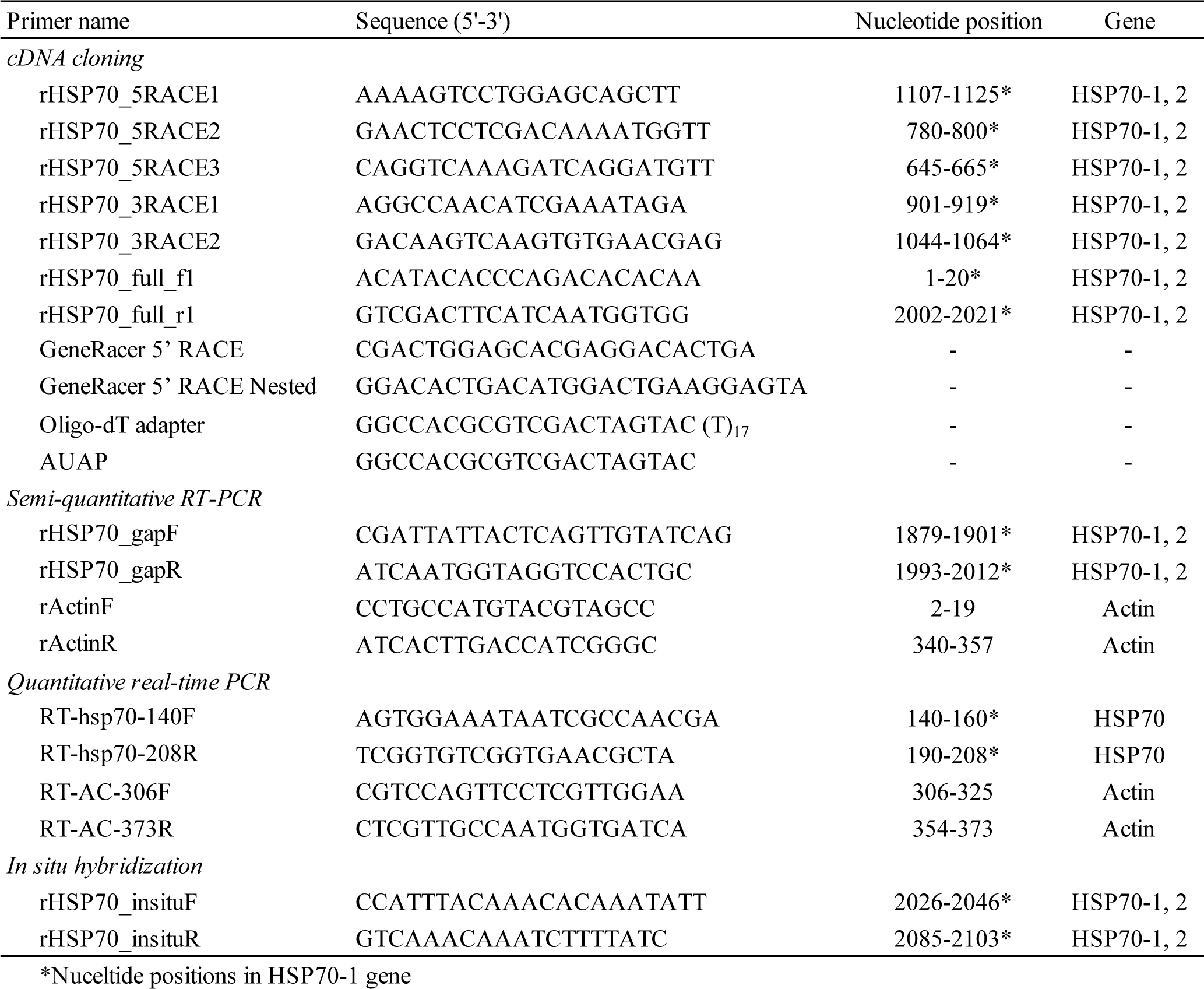
Nucleotide sequences of primers used in this study

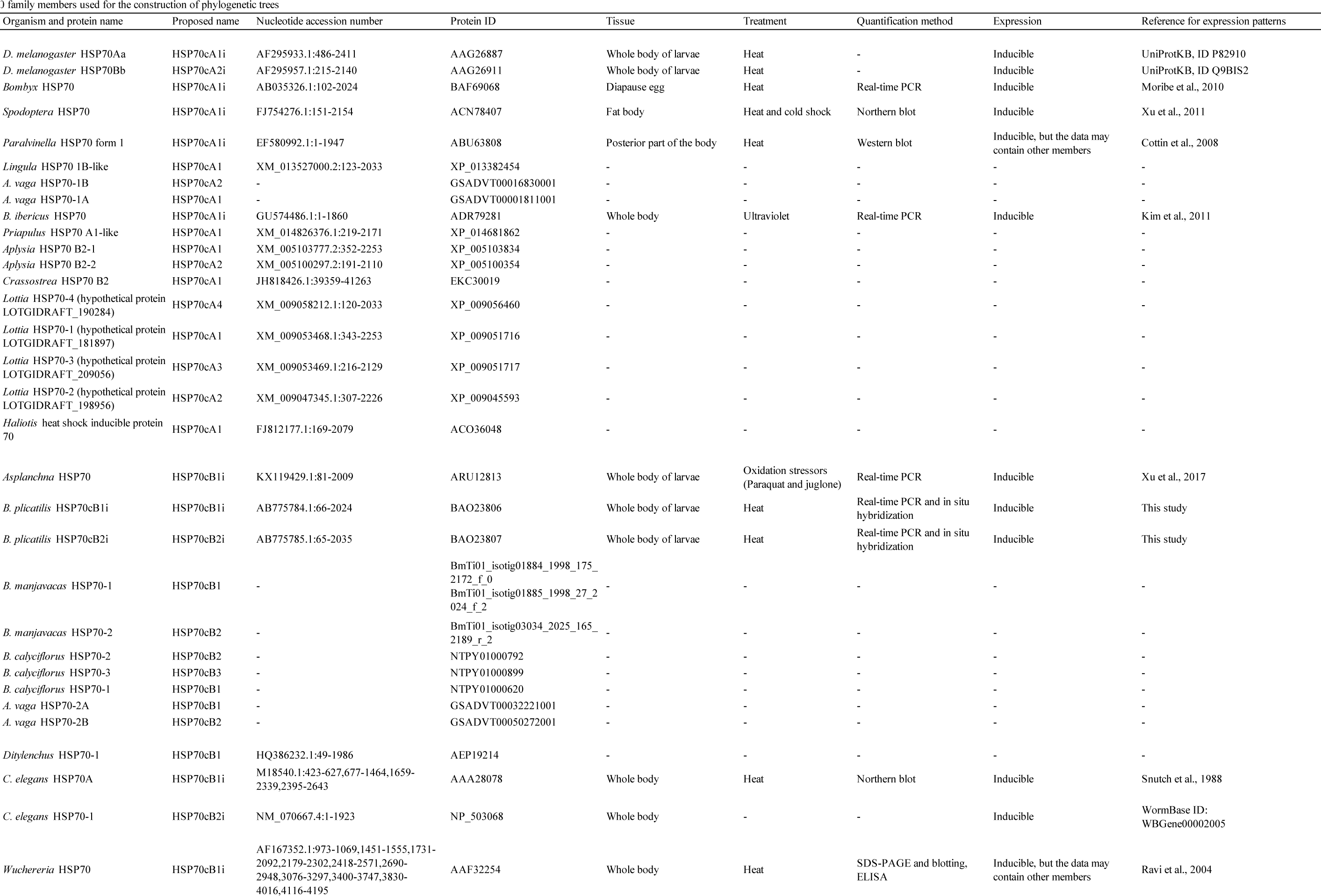
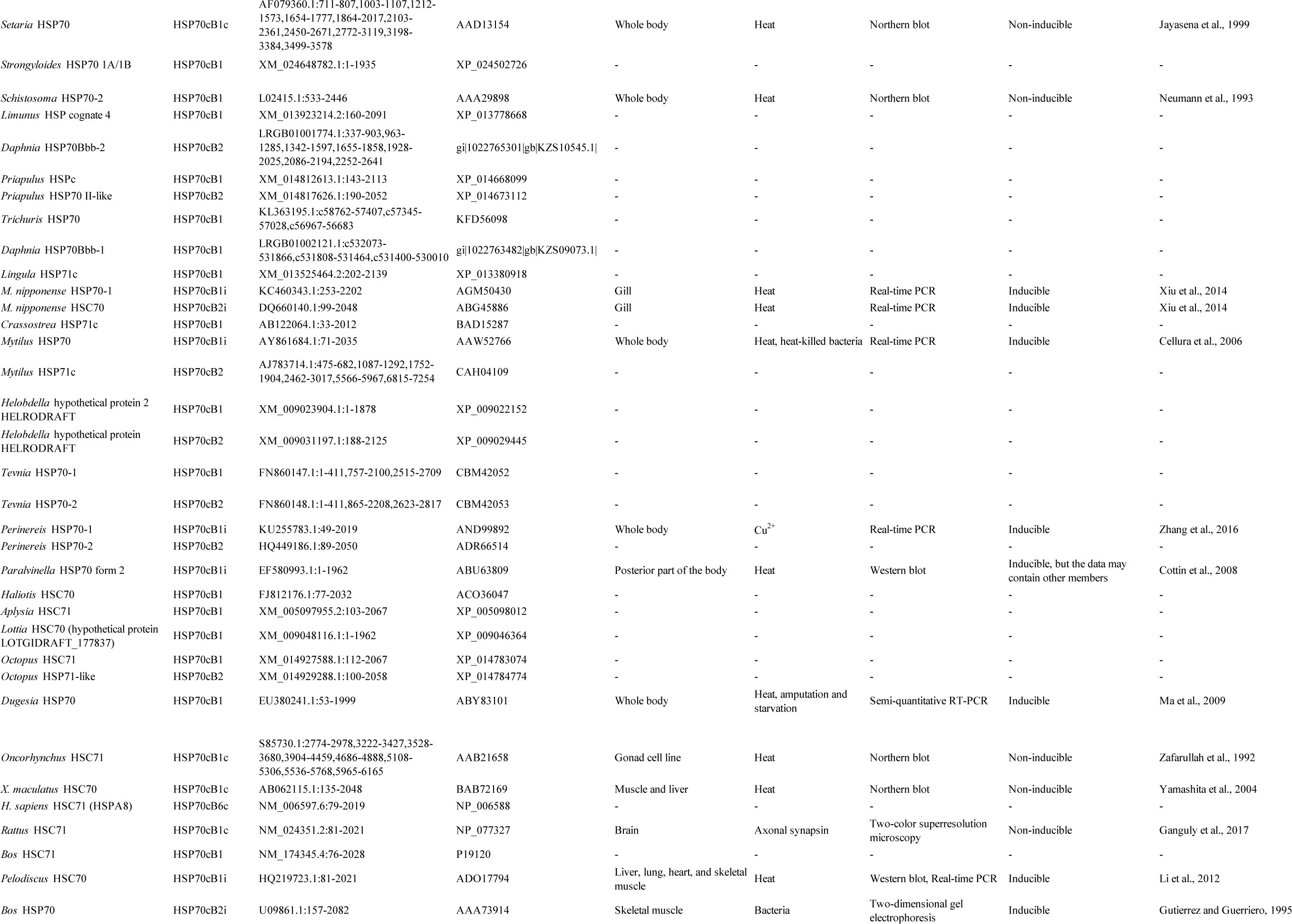
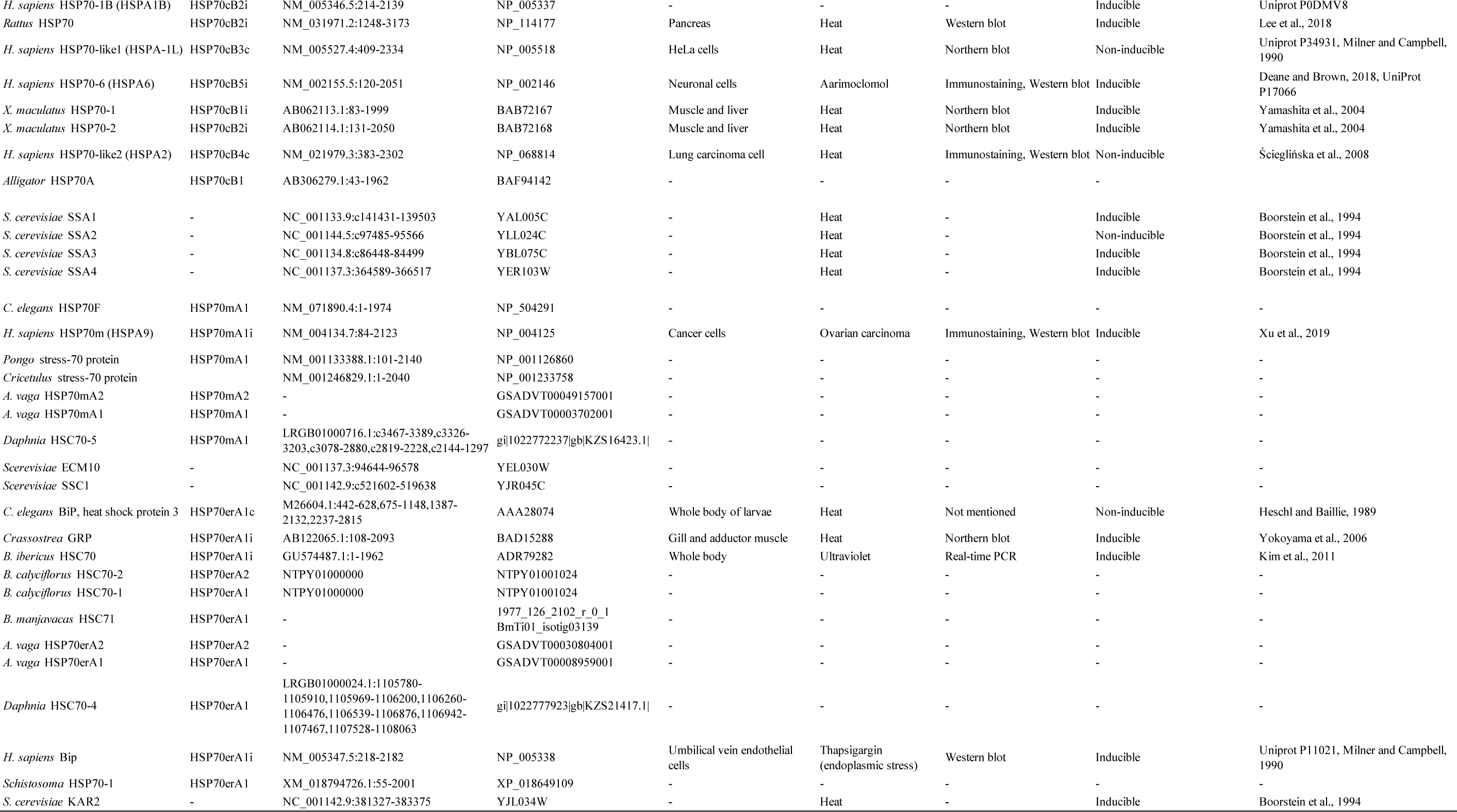

**Supplementary fig. S1.**
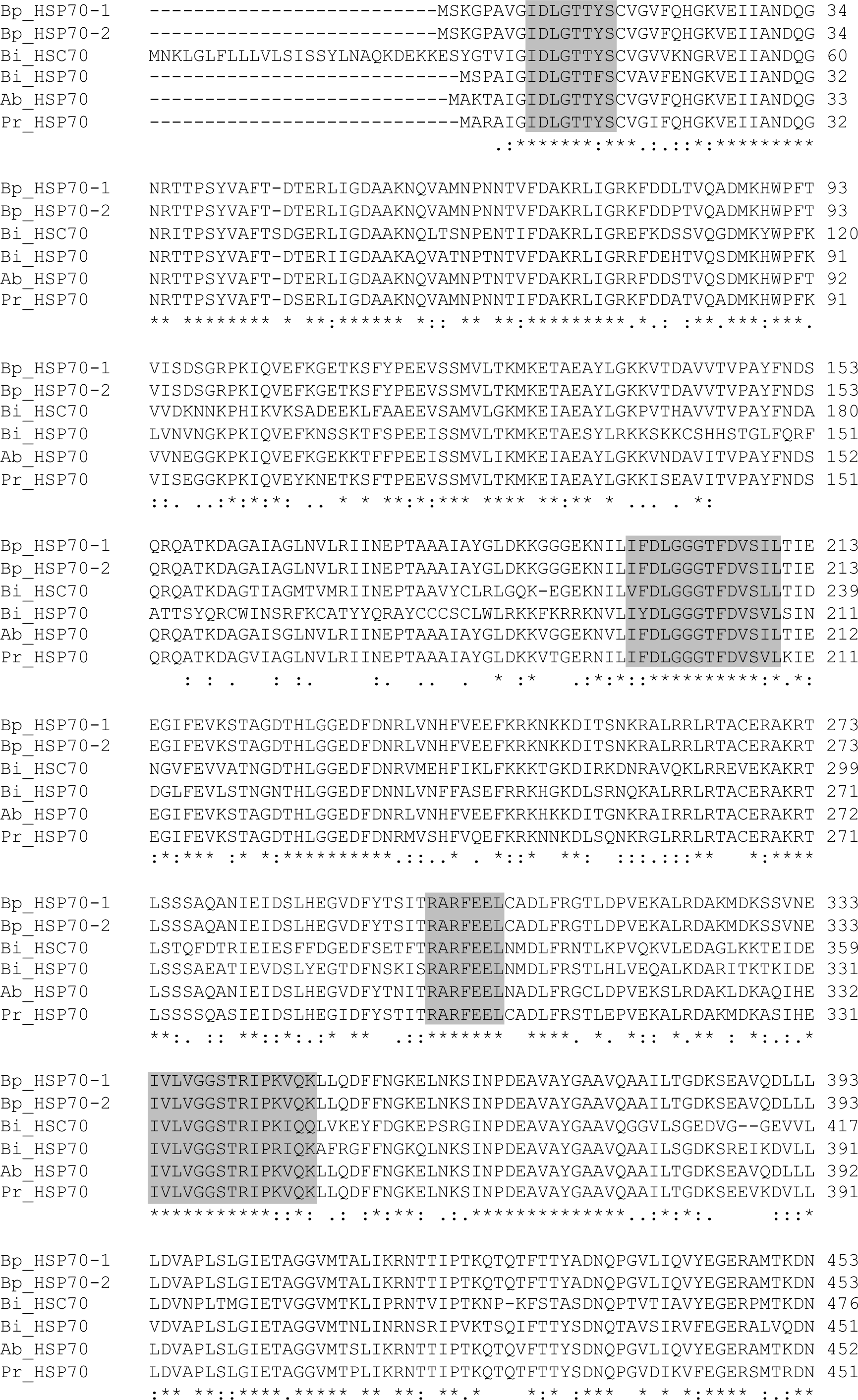
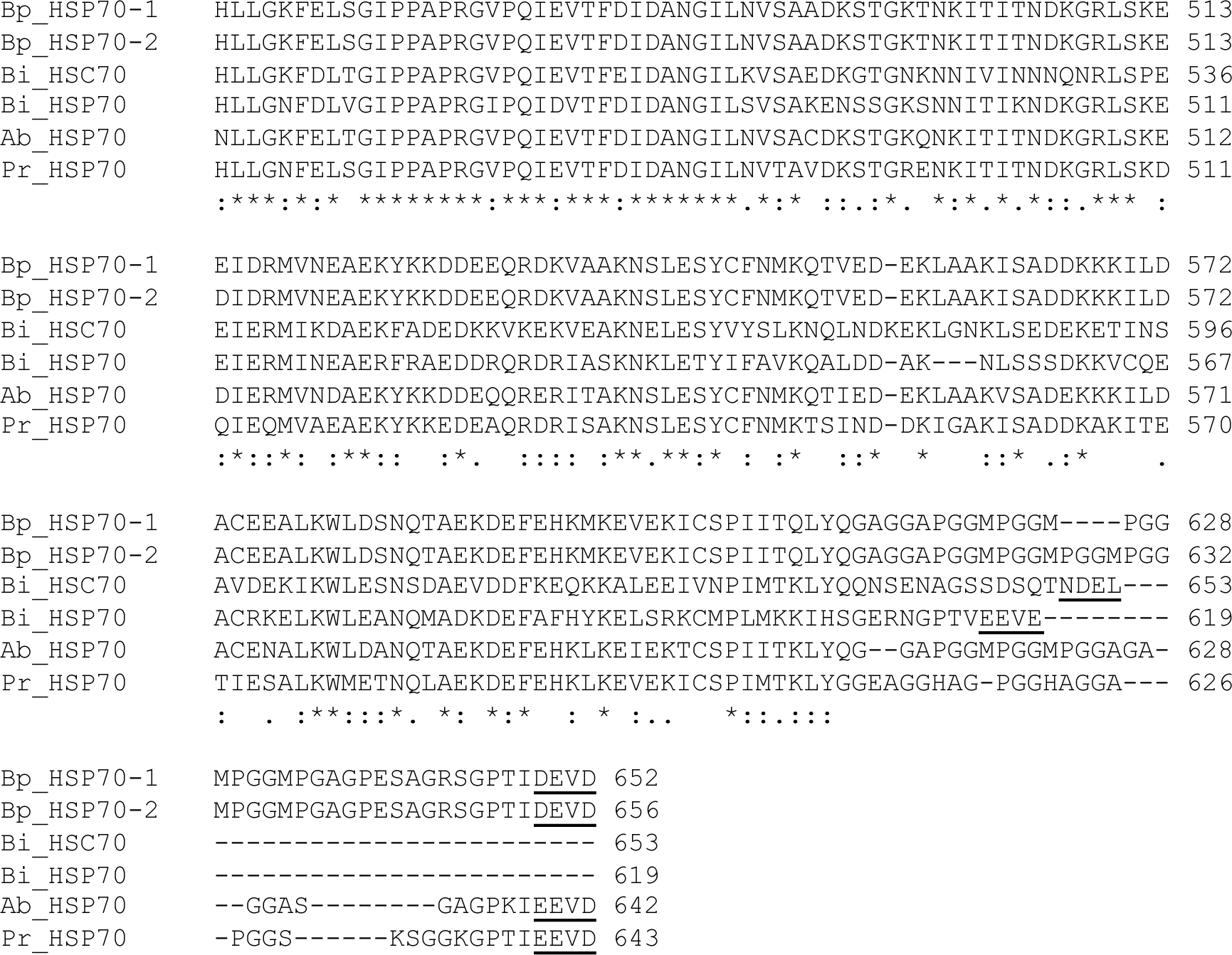
Comparison of the deduced amino acid sequences of cDNAs encoding HSP70-1 and HSP70-2 of *Brachionus plicatilis* with those of the HSP70 family members from other rotifers. Dashes denote gaps introduced to maximize homology. The HSP70 protein family signatures are meshed, and the EEVD motifs are boxed. The non-organelle consensus motifs and bipartite nuclear localization motifs are shown in blue and red letters, respectively. The numbers in the right margin of the sequences represent residues from the N-terminus. Bi, *B. ibericus* (ADR79281, ADR79282); Ab, *Asplanchna brightwelli* (ARU12813).

**Supplementary fig. S2.**
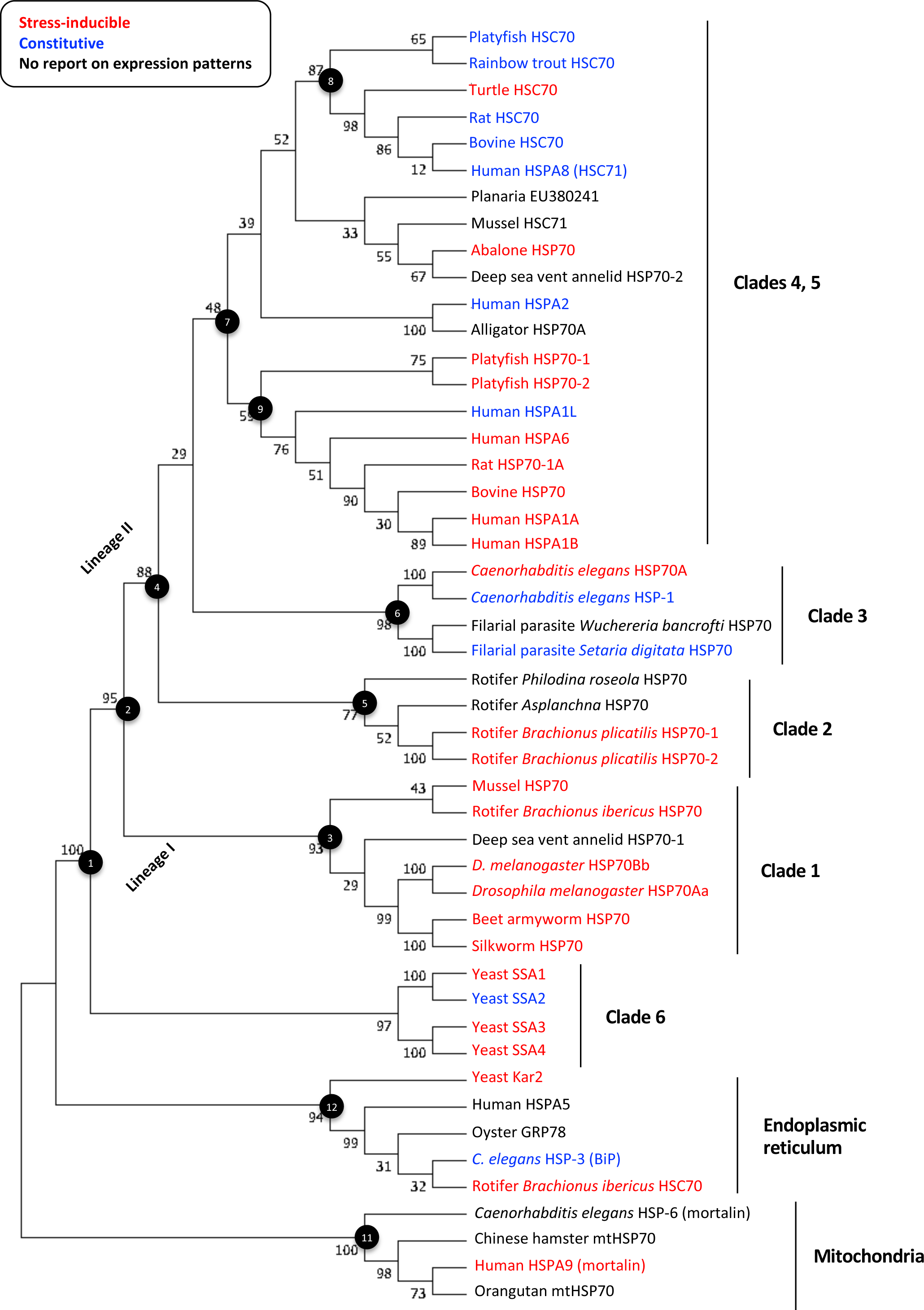
Maximum likelihood tree of HSP70 family members. A bootstrap consensus tree was constructed by the maximum likelihood method using the LG + G model. The bootstrap values from a 1000-replicate analysis are given at the nodes in percentage. Nodes and clades in fig. 2 are indicated in the tree. Stress-inducible and constitutive genes are shown in red and blue, respectively. No data on the stress inducibility is available for genes shown in black.

**Supplementary fig. S3.**
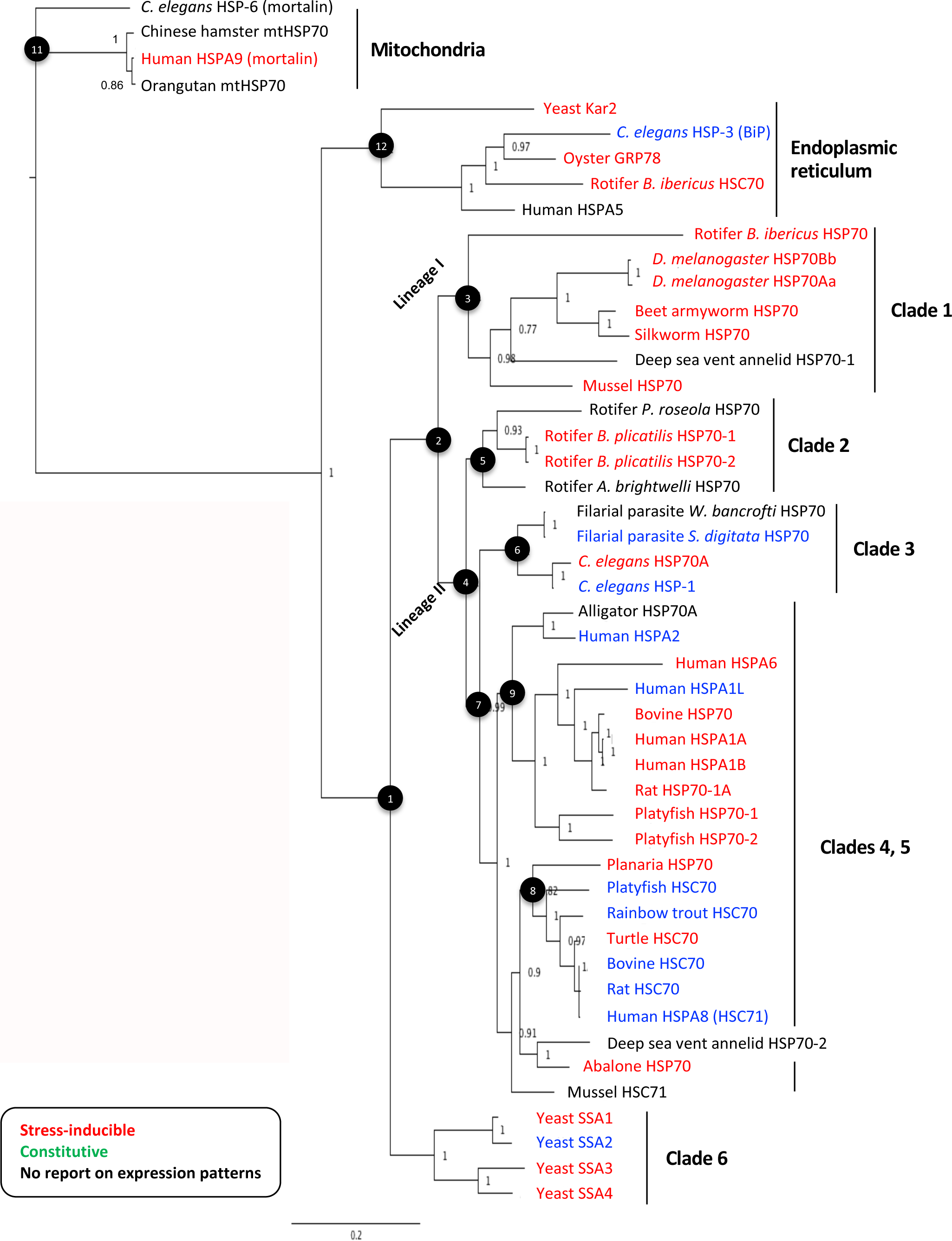
Bayesian consensus tree of HSP70 family members. Nodes and clades in fig. 2 are indicated in the tree. Numbers on the branches indicate the posterior probability. Stress-inducible and constitutive genes are shown in red and blue, respectively. No data is available on the stress inducibility for genes shown in black.

**Supplementary fig. S4.**
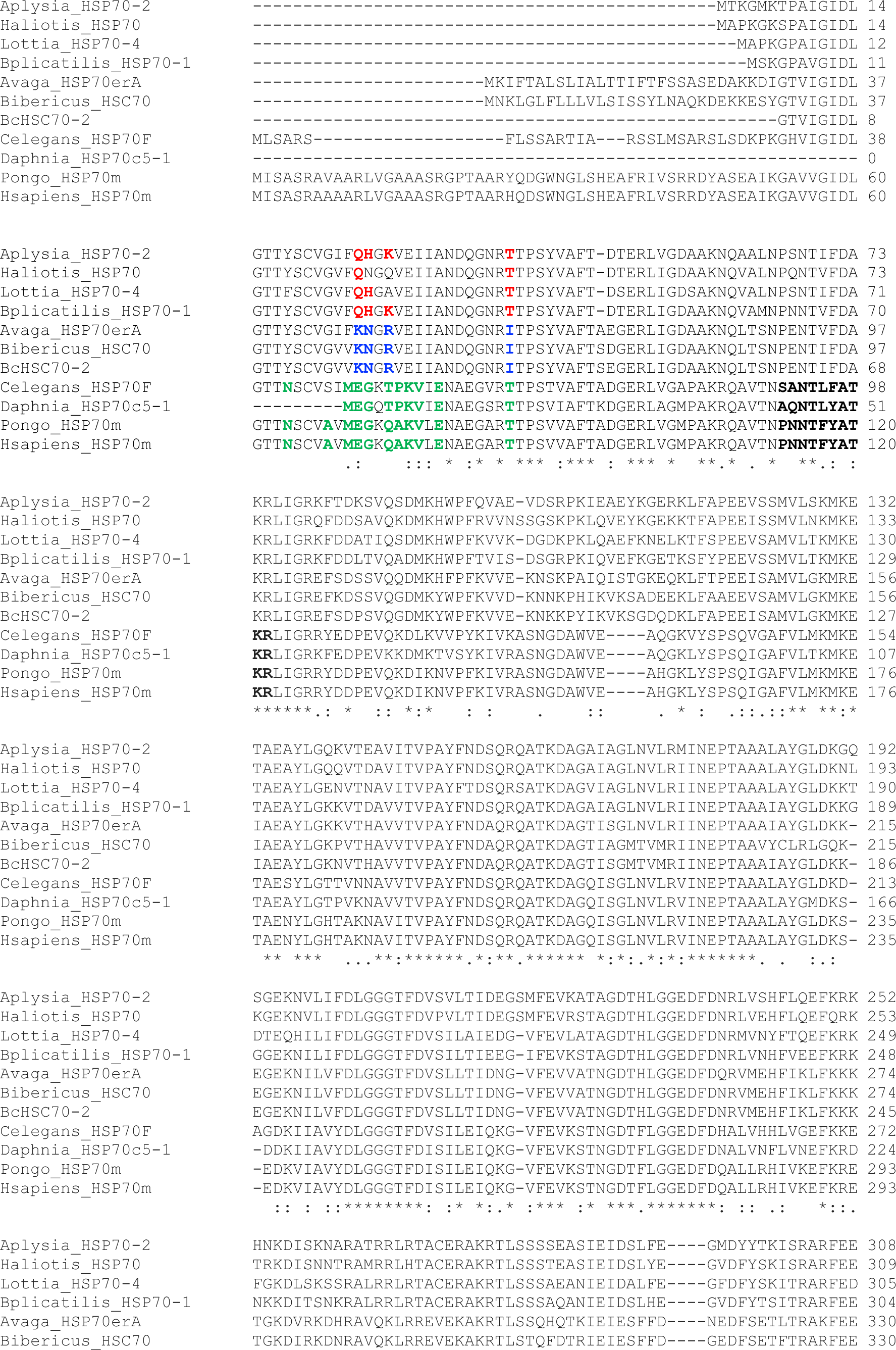
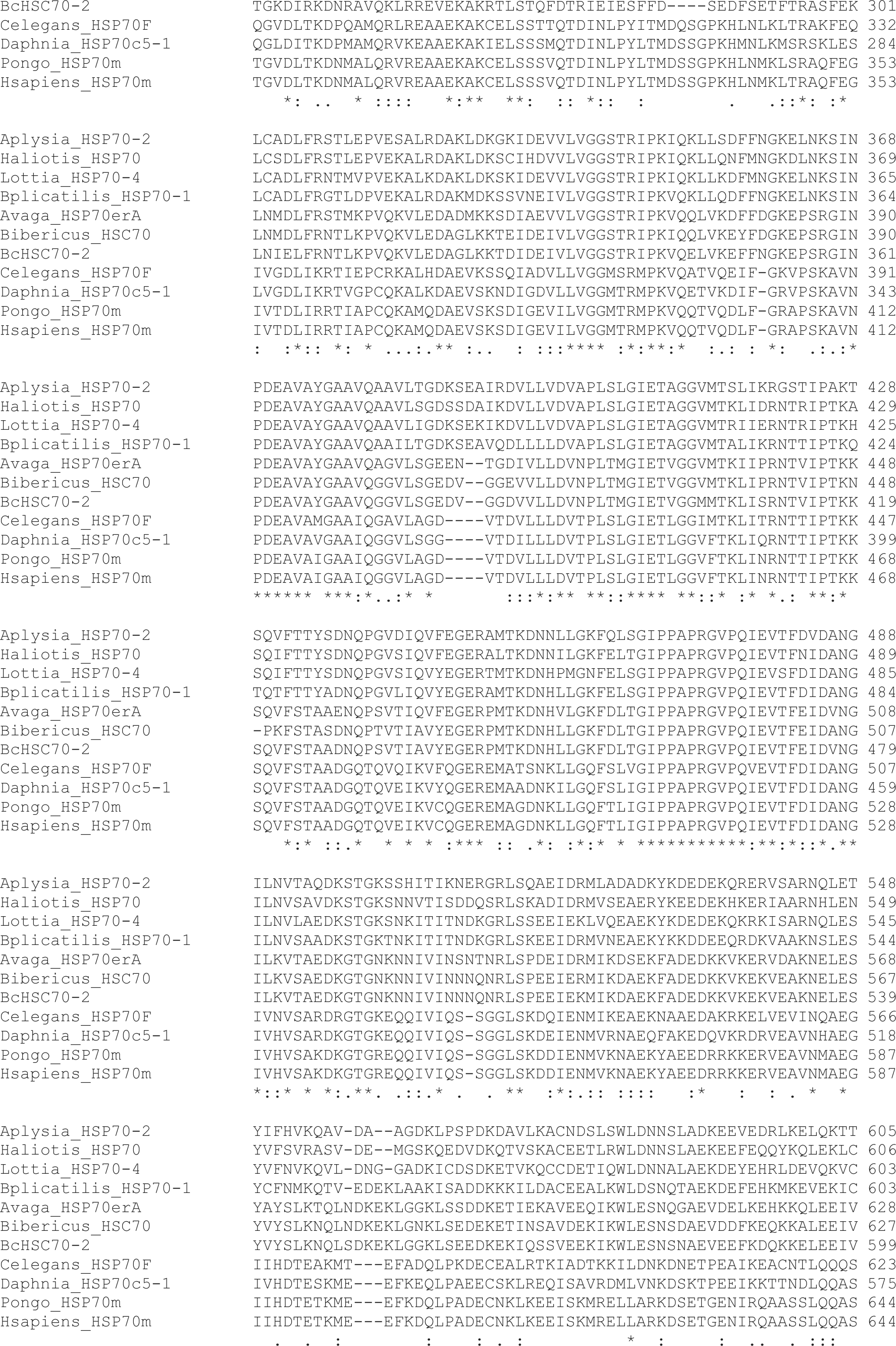
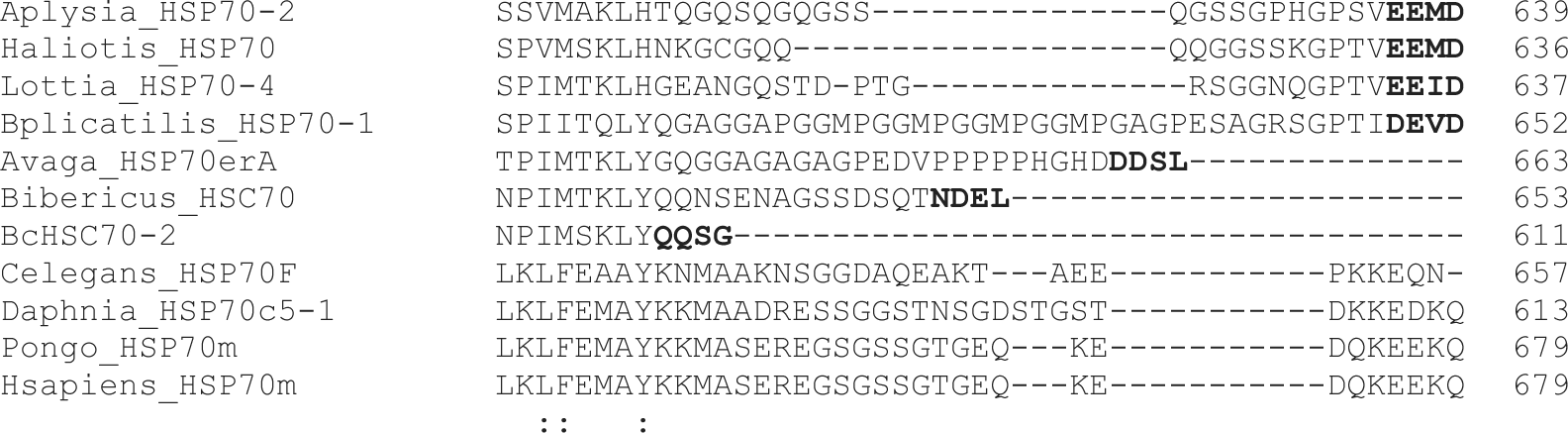
Alignment of several HSP70 family members with variations in the cytosolic (EEVD), endoplasmic reticulum (ER; HDEL/KDEL), and mitochondrial (PEAEYEEAKK) motifs. The three motifs are shown in bold. Distinctive amino acid residues in the newly identified motifs are shown in red (cytosolic), blue (ER), and green (mitochondrial). The discrimination of organelle-specific HSP70 family members becomes more evident by using multiple motifs.

**Supplementary fig. S5.**
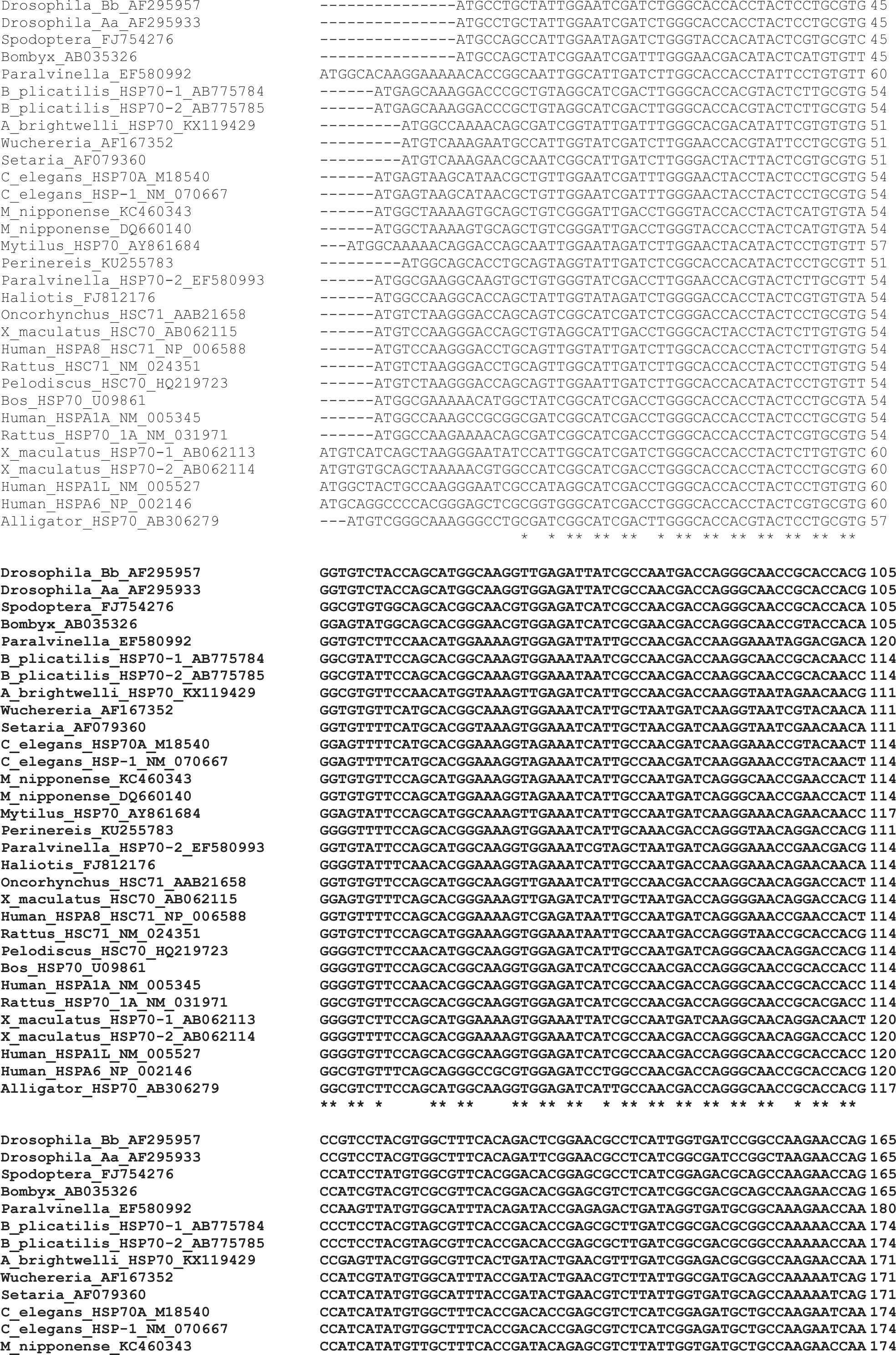
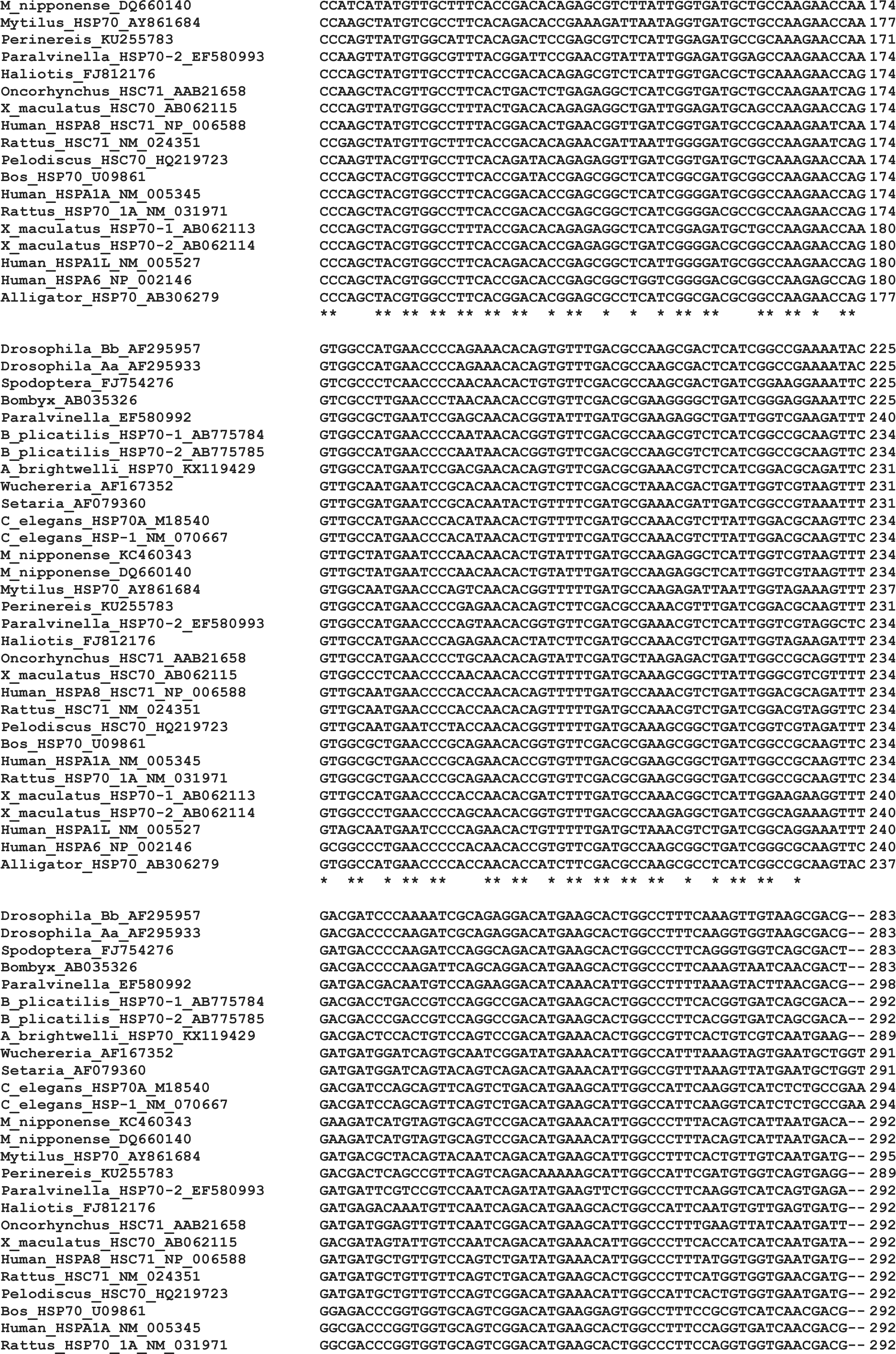
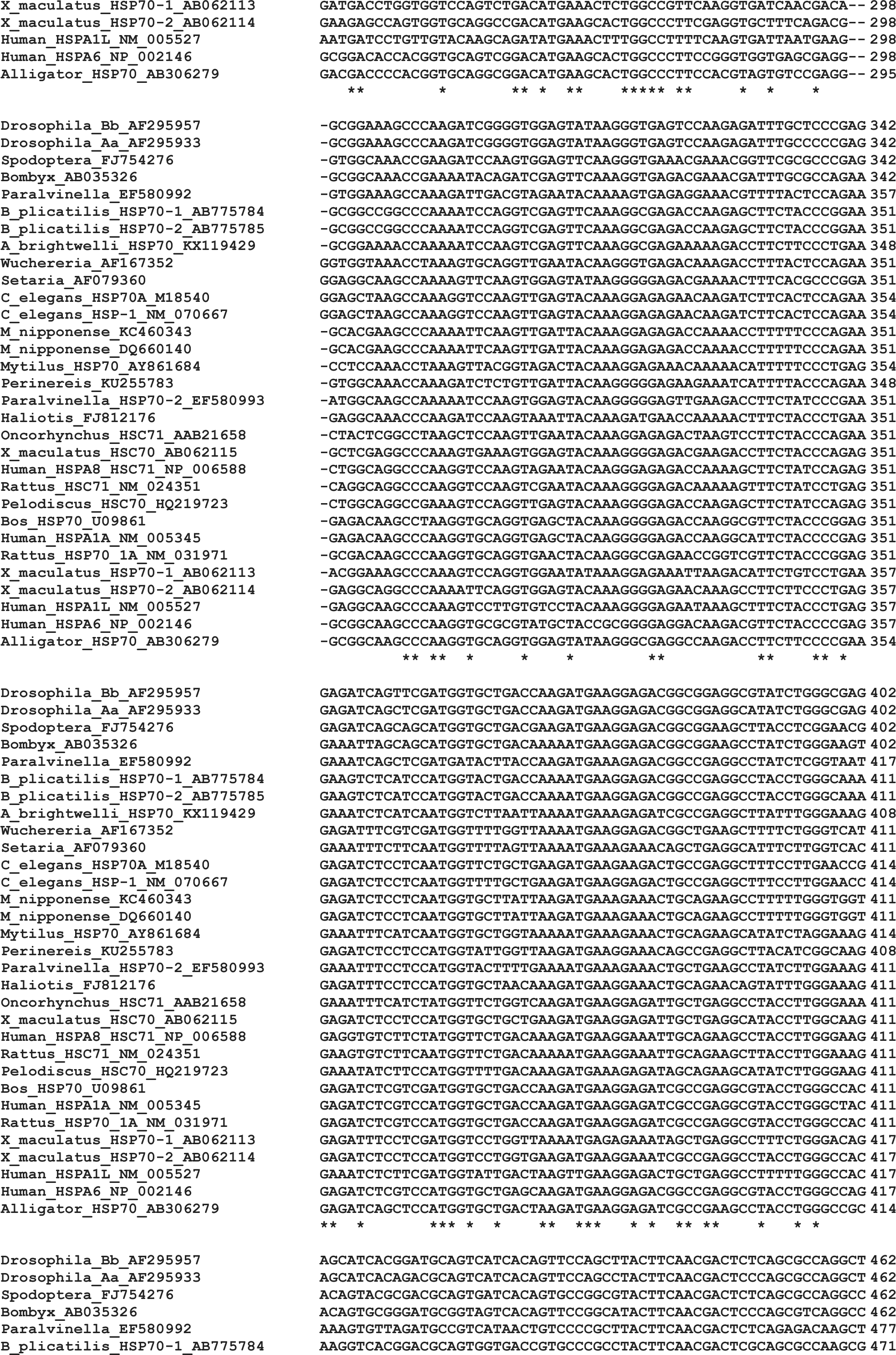
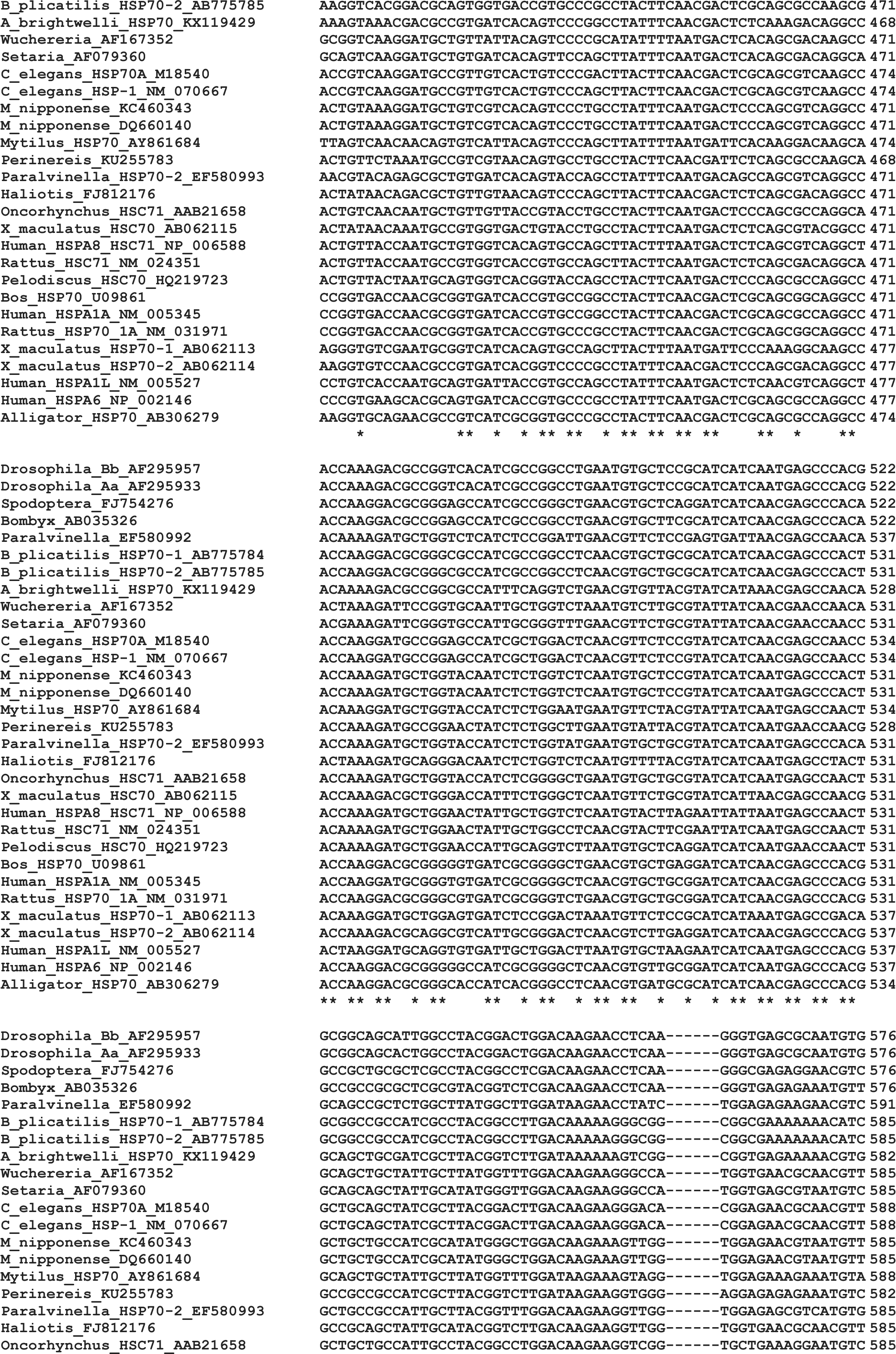
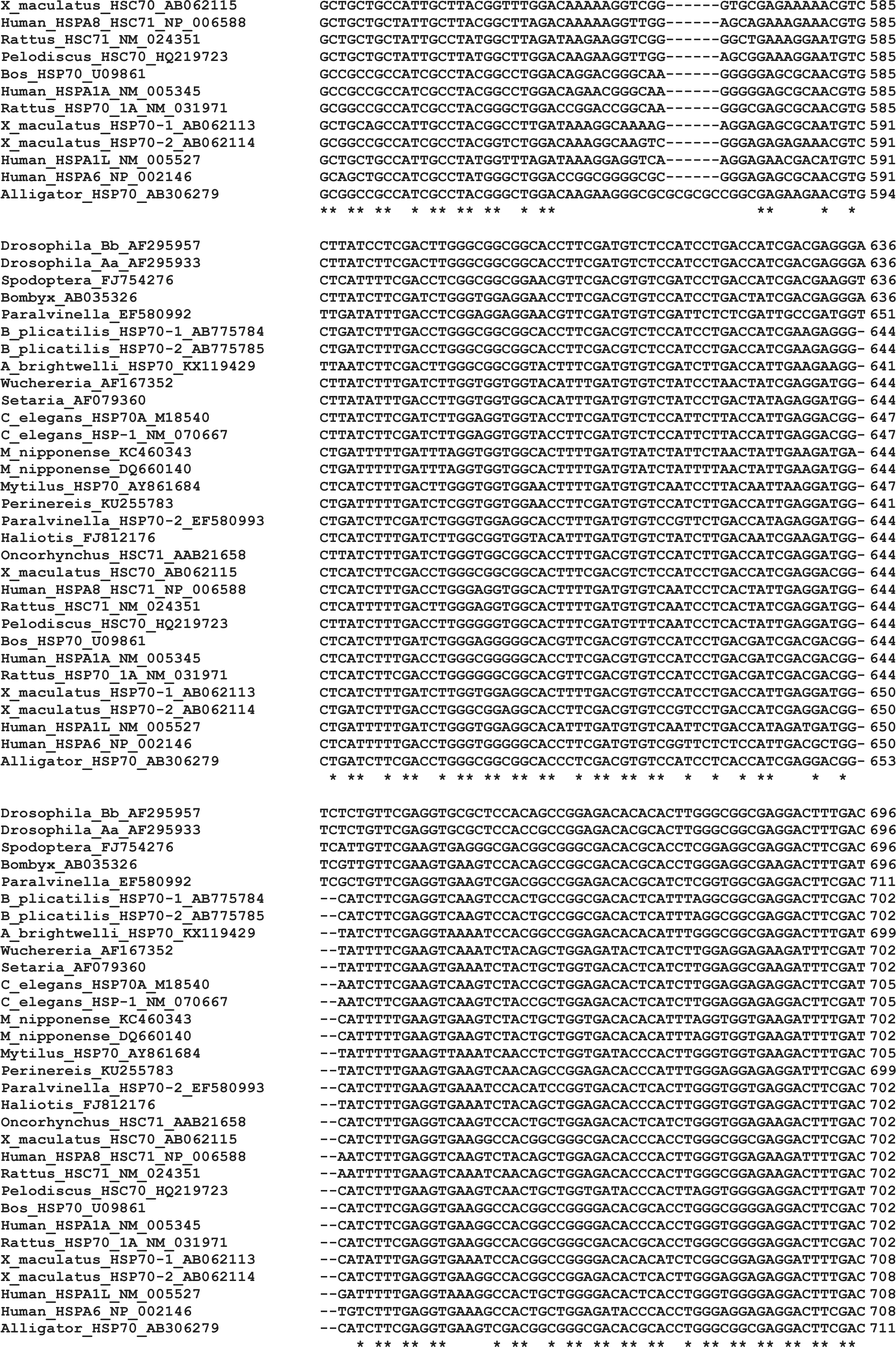
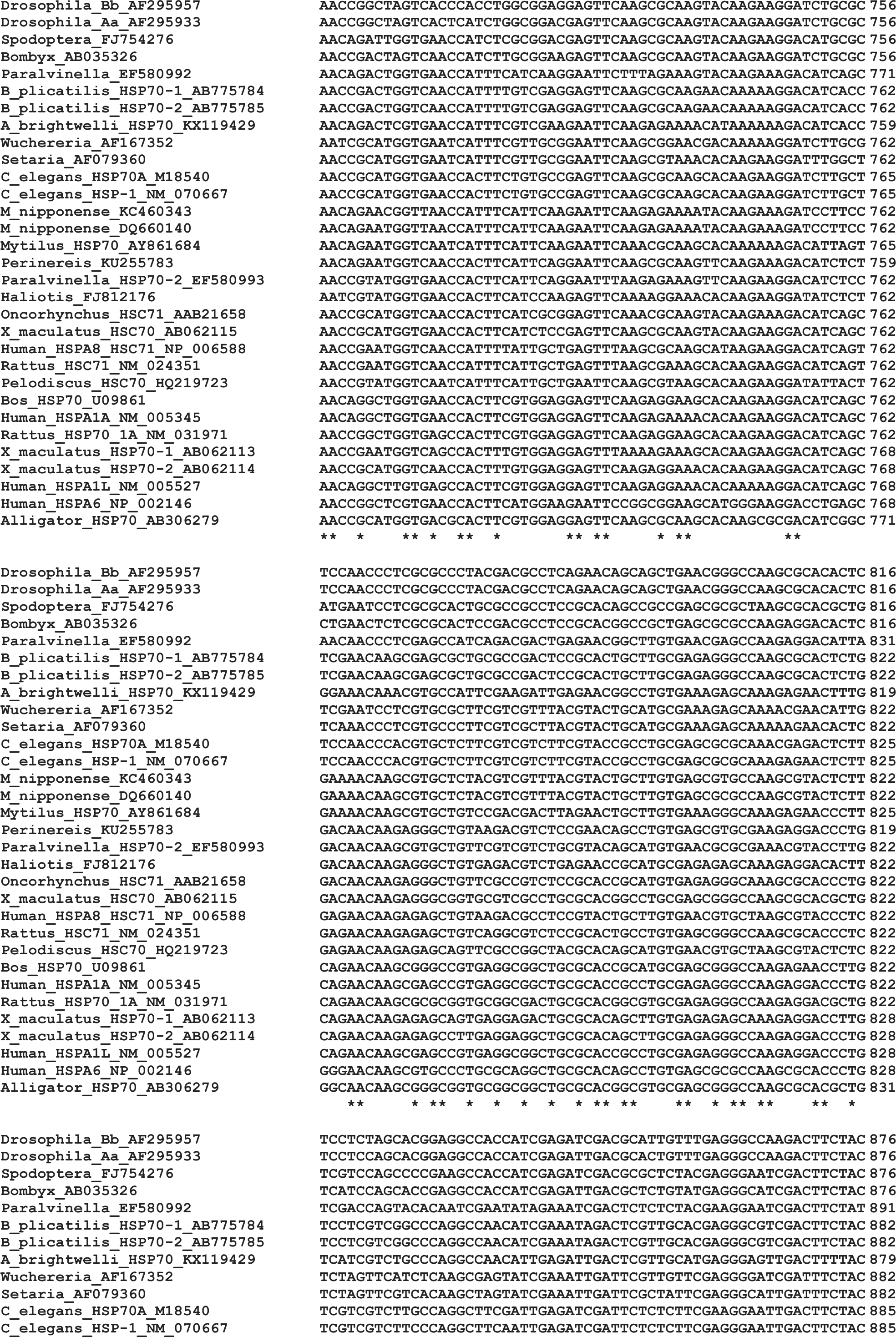
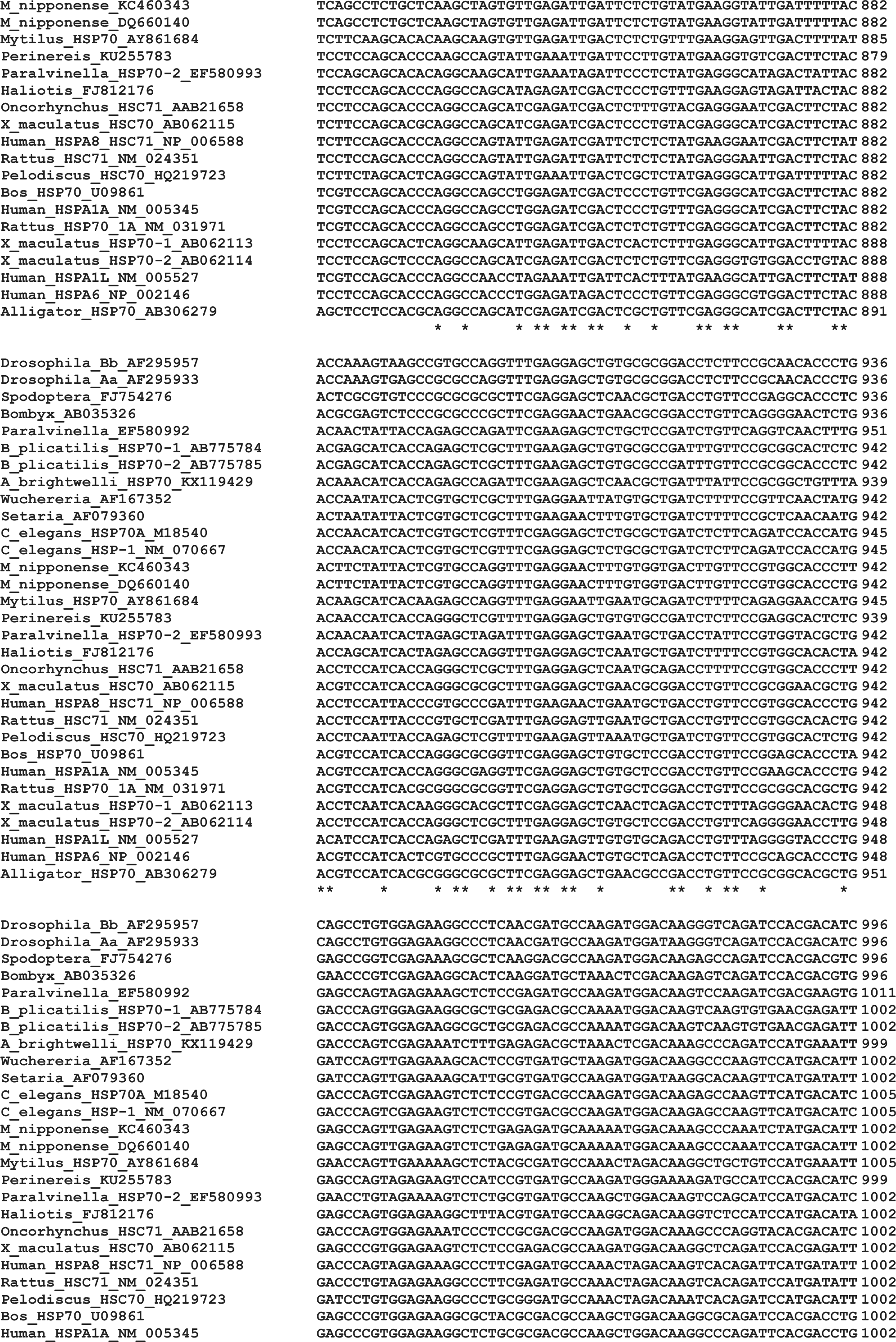
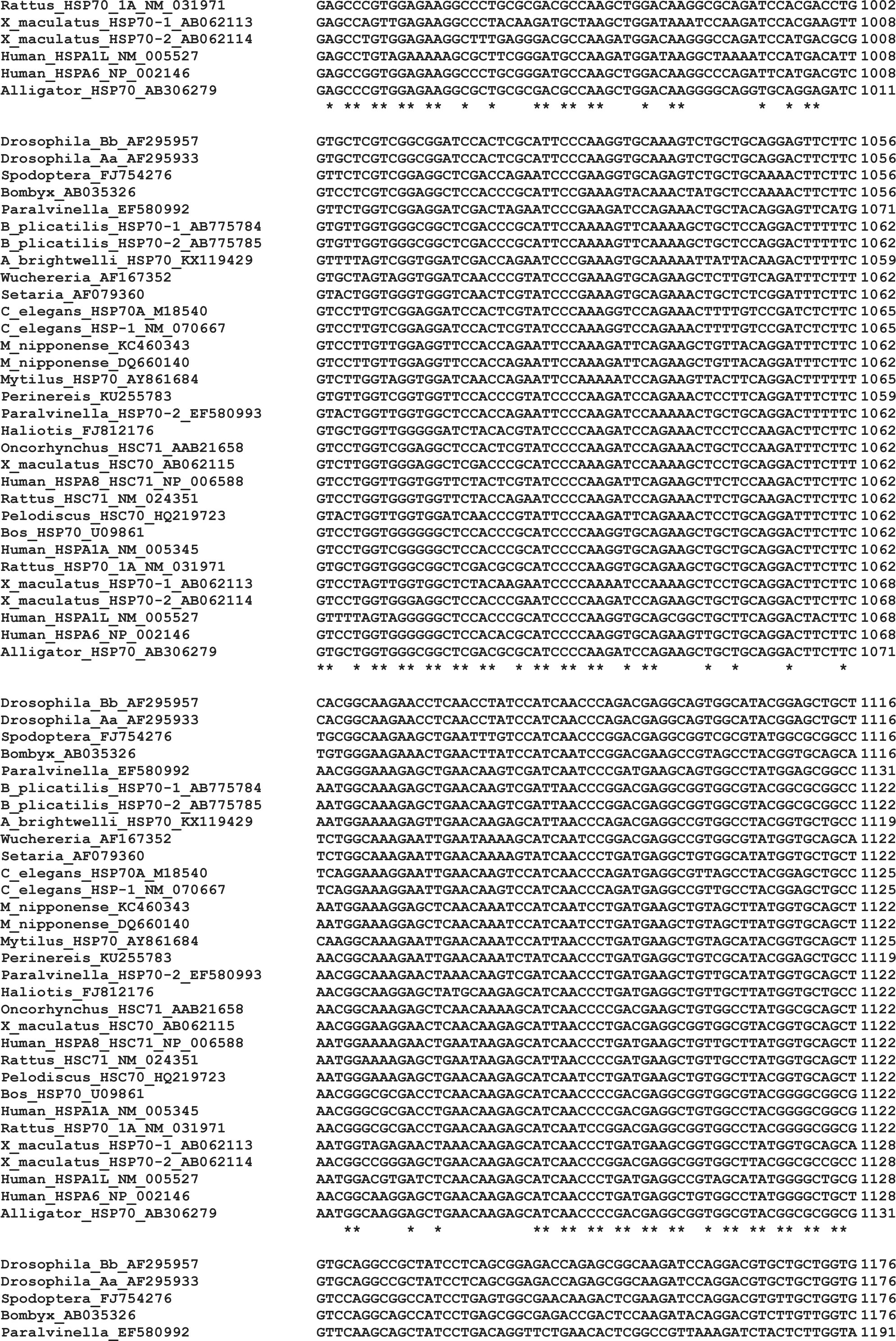
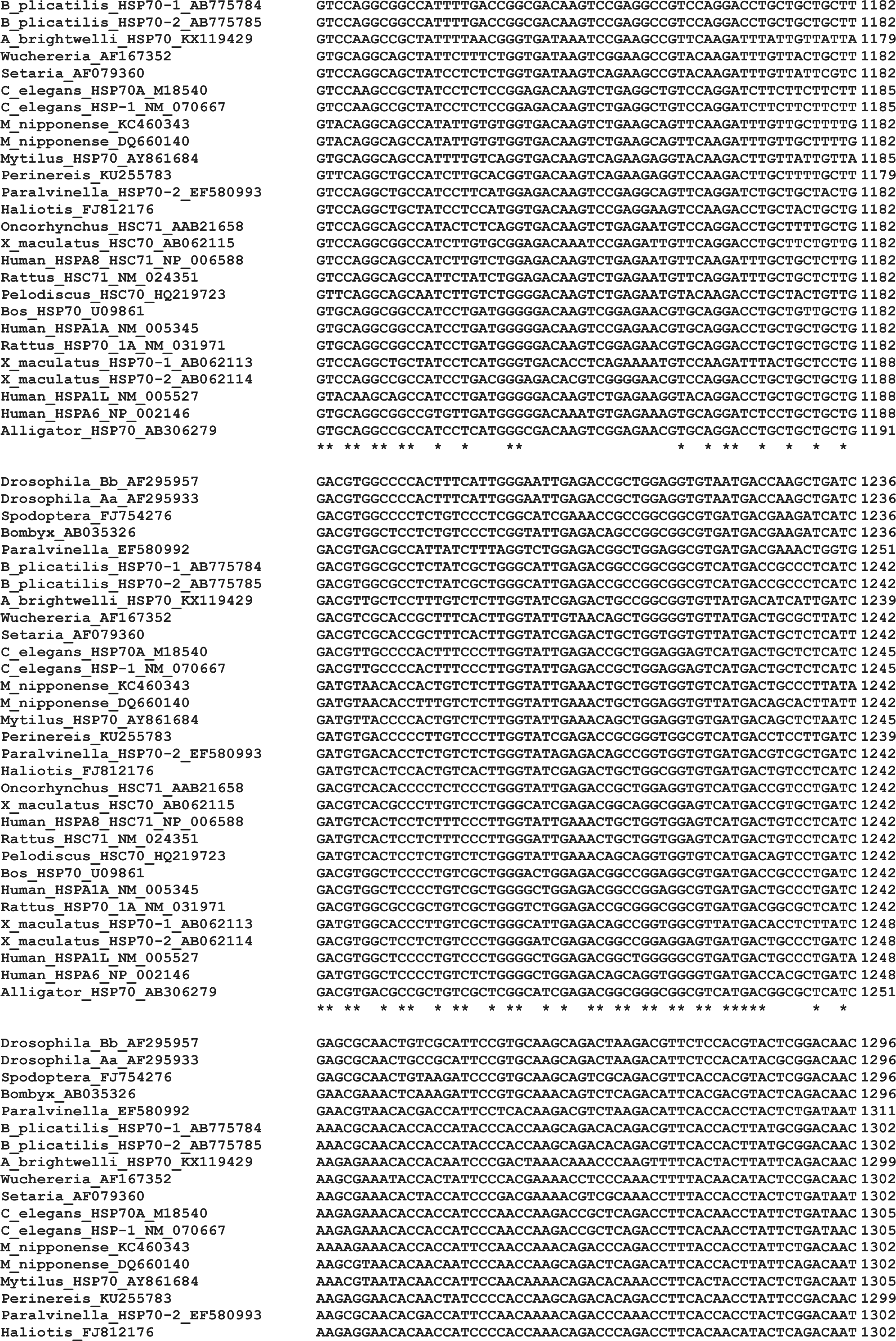
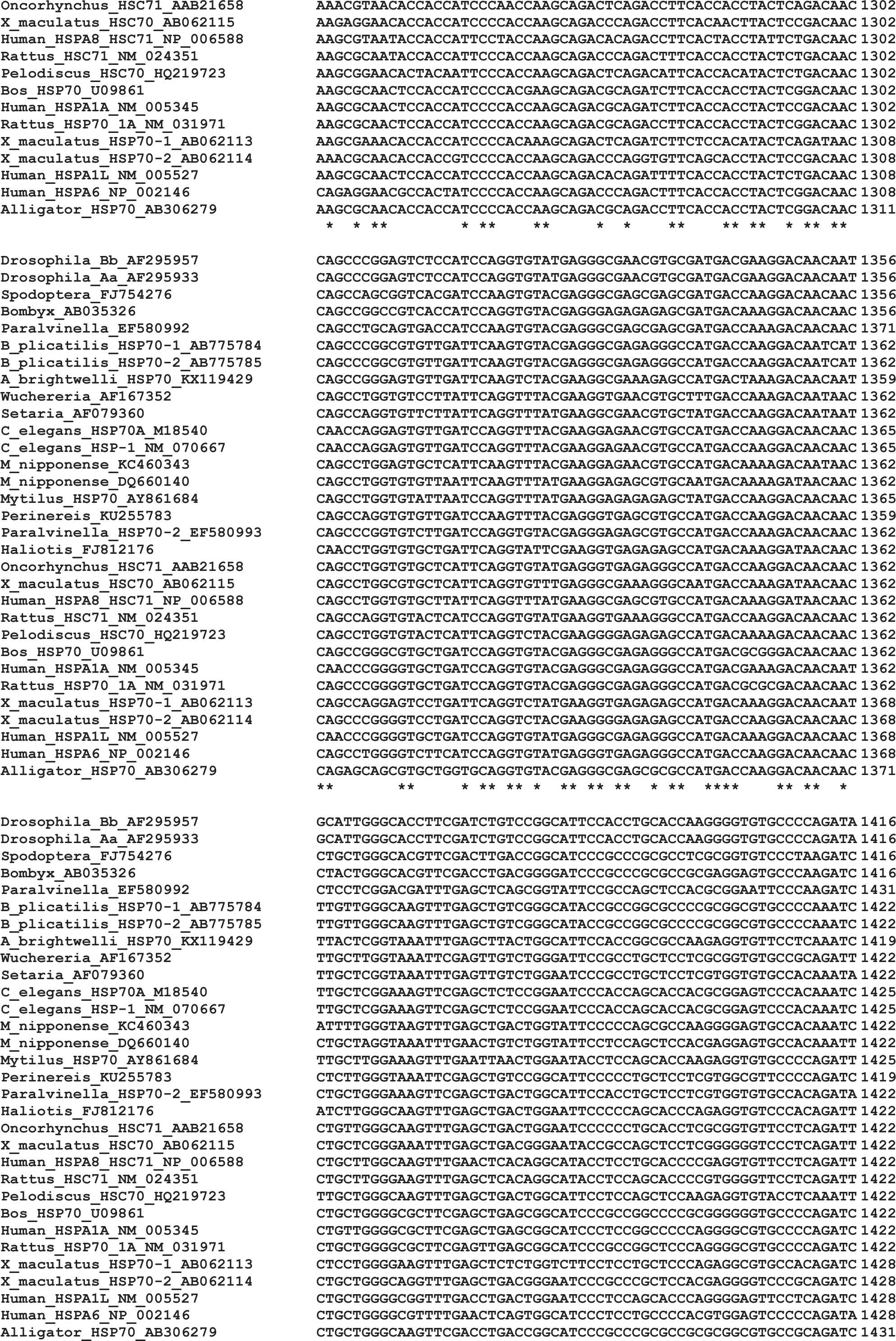
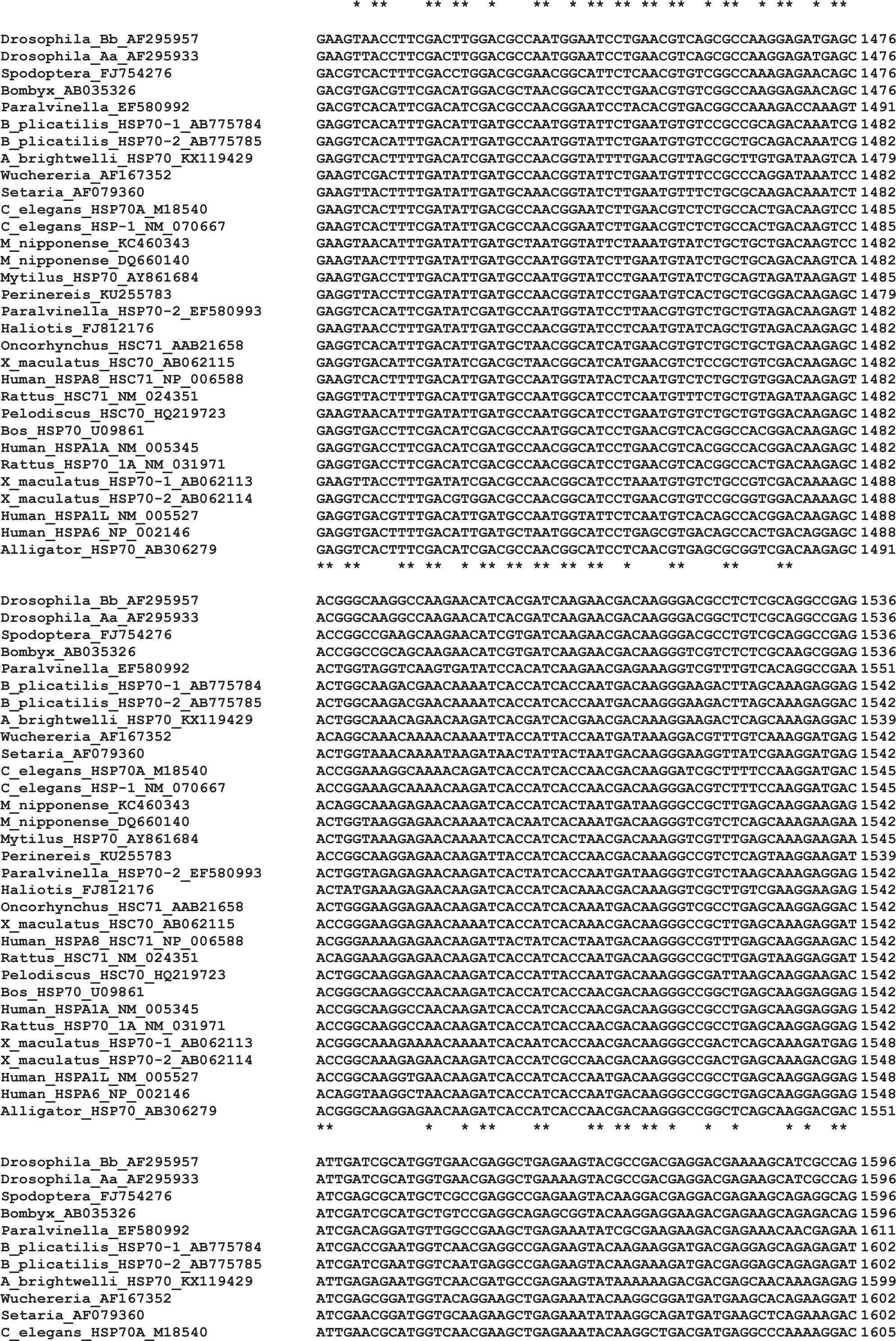
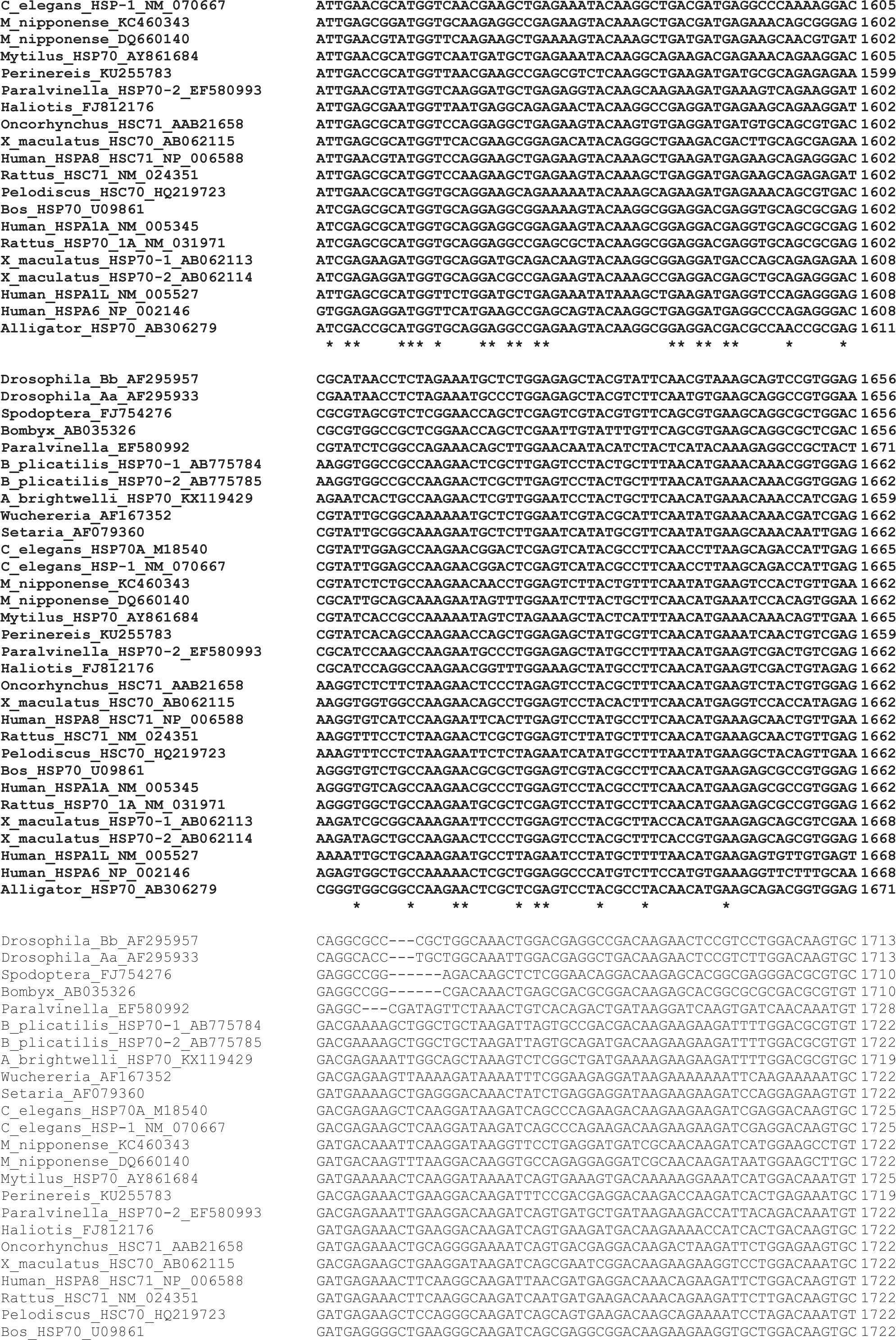
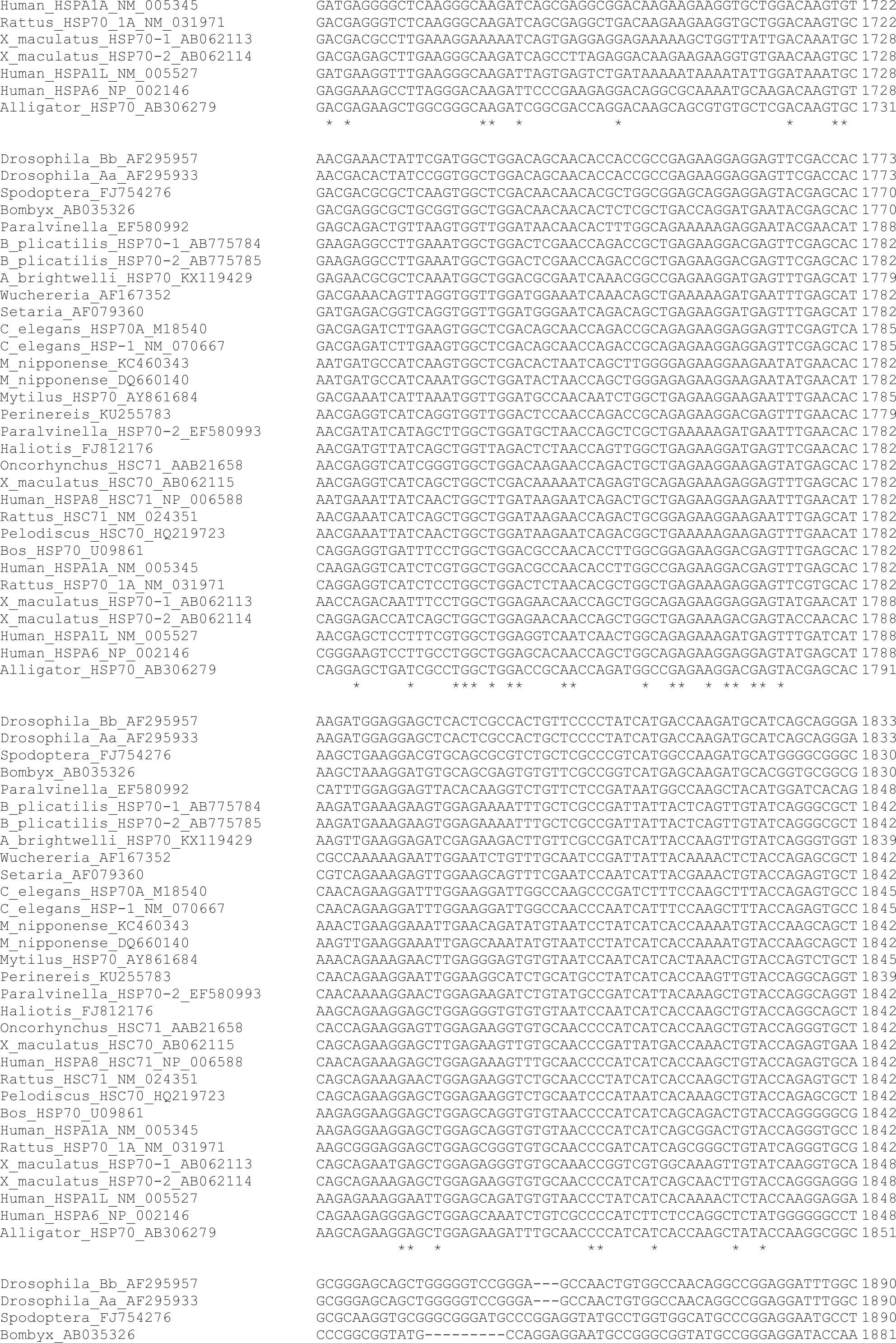
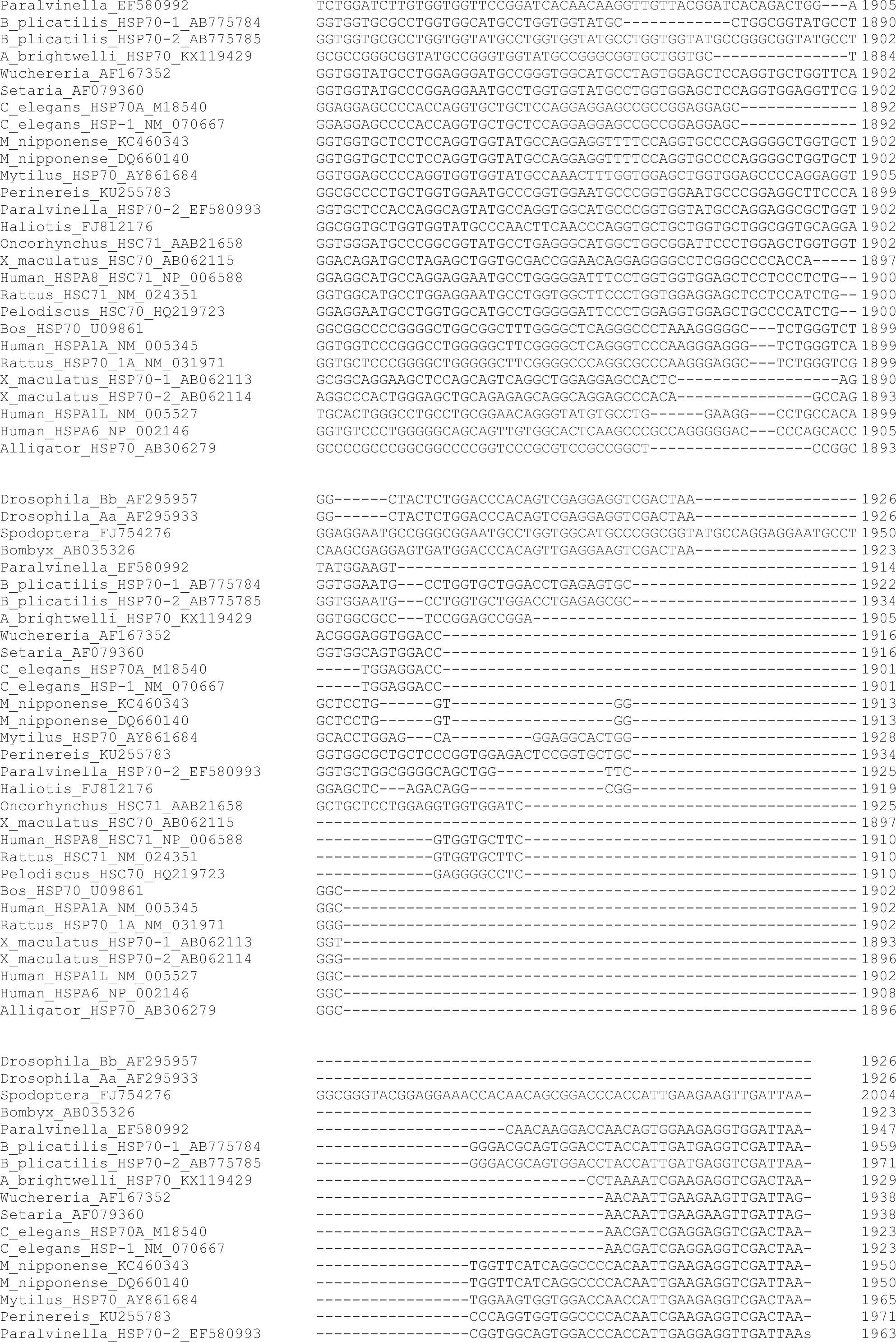
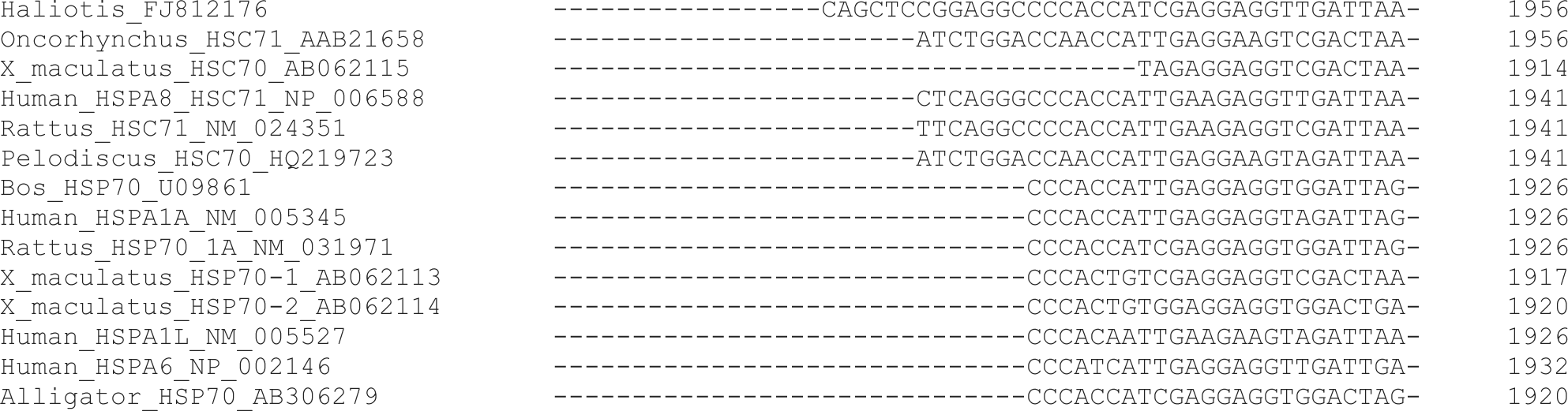
Comparison of HSP70 nucleotide sequences used for the calculation of synonymous and nonsynonymous substitution rates. Regions shown in bold were used for the calculation.

**Supplementary fig. S6.**
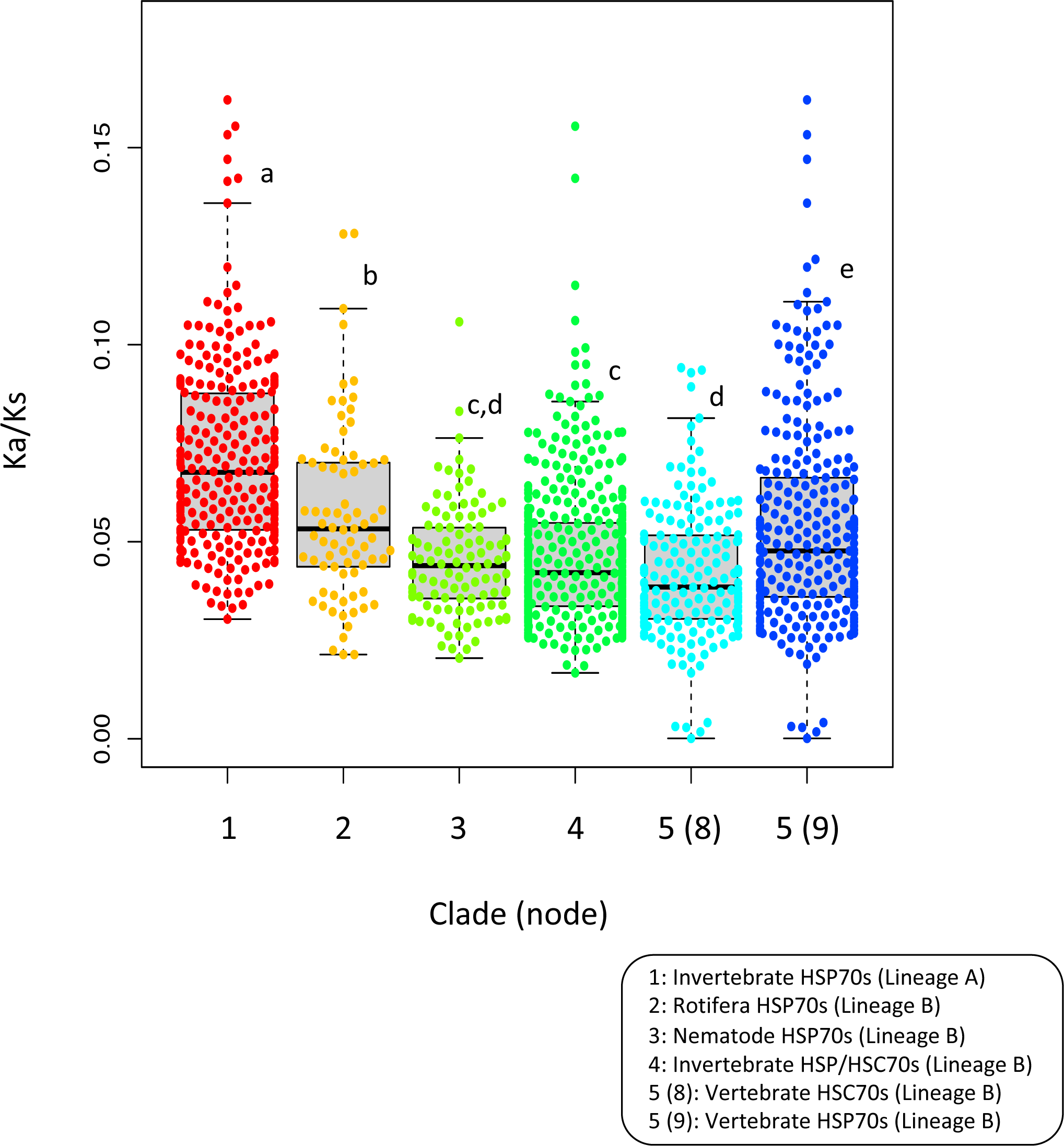
Bee swarm boxplots of Ka/Ks values for each clade. Only Ka/Ks values calculated from inter-cluster pairs were averaged. Statistical differences were calculated by Kruskal-Wallis test (chi-squared = 200.45, df = 5, p-value < 2.2e-16) followed by the non- parametric post-hoc tests (pairwise Wilcox test with P value adjustment by the Holm method). Clades sharing same letters are not significantly different at the 5% level of significance. One outlier in clade 3 (0.53) is not included in the plot although this value was used for all statistical analyses.

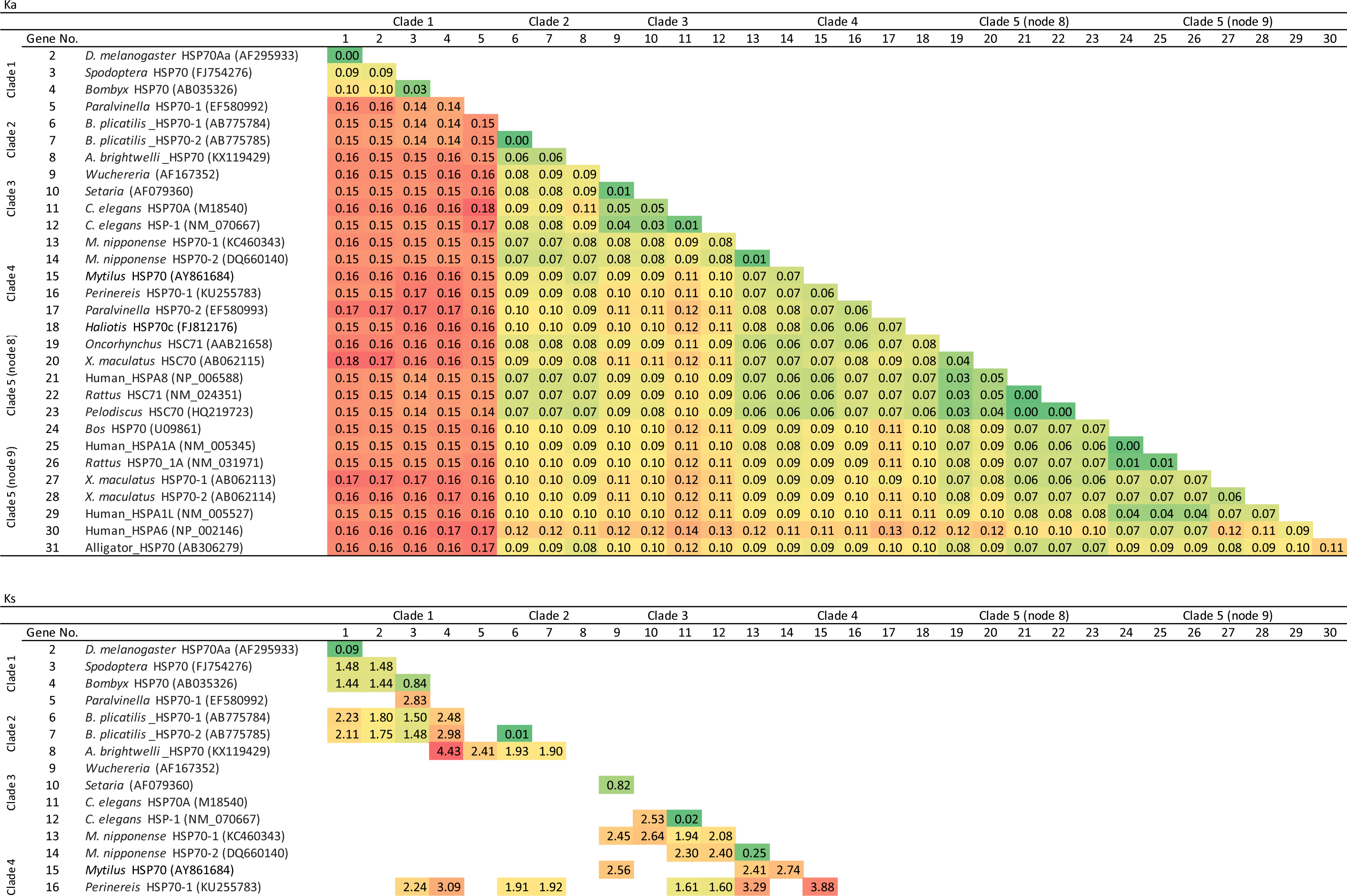
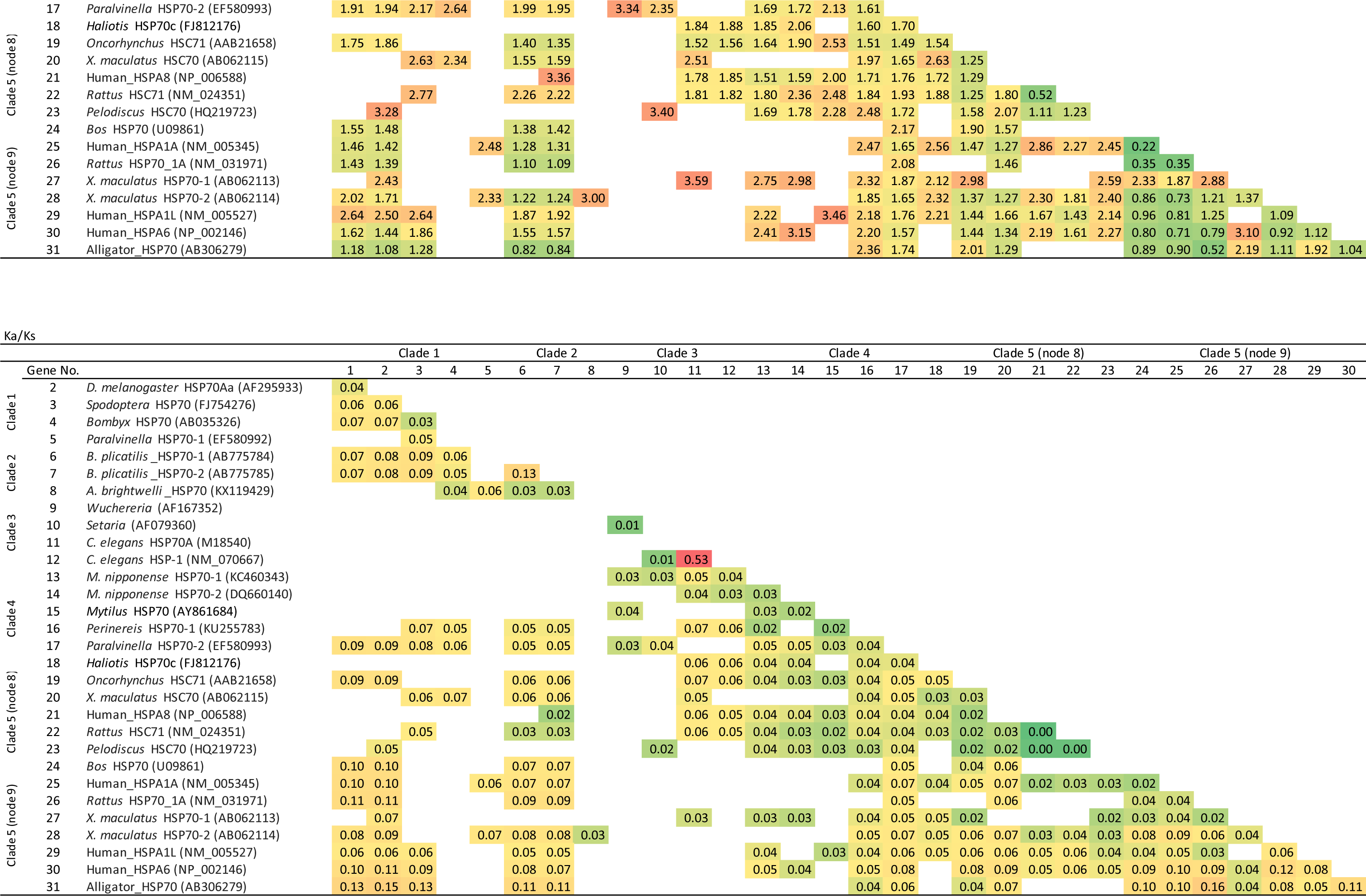

## Supplementary material 2

### Heat Treatment

For semi-quantitative RT-PCR, rotifers were batch-cultured in 400 mL at 25°;C. In the exponential growth phase, approximately 6 × 10^4^ individuals were randomly divided into two groups and concentrated to about 10 mL by filtration through 20 µm mesh filters. One group (heat-treated group) was transferred to culture medium at 30°;C for 10 min, collected with a 20 µm mesh filter, and stored in ISOGEN (Nippon Gene, Tokyo, Japan) at -80°;C. The control group was prepared by the same procedure at 25°;C.

For quantitative real-time PCR, 8 rotifers bearing 2 or 3 eggs were selected from a batch culture population and inoculated into a well of a 12-well plate (Nunc, Rochester, NY) containing 1 mL of culture medium at 25°;C. The 12-well plate was put on a water bath at 40°;C for 10 min, and then transferred to a water bath at 25°;C. Rotifers were sampled before heat treatment (control), just after heat treatment (time 0), and 2, 3, 4, 6, and 8 h after heat treatment (n = 3, each group contained 8 individuals). Rotifer samples were stored in 500 µL ISOGEN (Nippon Gene) and stored at -80°;C until use. Rotifers used for *in situ* hybridization were prepared by the same method except the heat shock temperature was 30°;C.

### cDNA Cloning of HSP70

5’ RACE for the partial sequence of *B. plicatilis* HSP70 gene [AB076052; (Kaneko et al. 2002)] was carried out using a GeneRacer kit (Invitrogen, Carlsbad, CA) with gene-specific primers rHSP70_5RACE1, rHSP70_5RACE2, and rHSP70_5RACE3 (supplementary table S1). Total RNA was extracted from a rotifer population in an exponential growth phase and used for 1st strand cDNA synthesis using the rHSP70_5RACE1 primer and the GeneRacer kit (Invitrogen). First round PCR was performed at 94°;C for 3 min followed by 30 cycles of 94°;C for 30 s, 55°;C for 30 s and 72 °;C for 1 min. The final extension step was performed at 72°;C for 5 min. The 20 µL-reaction mixture contained about 1 µg of the 1st strand cDNA, 2 µL of Ex *Taq* buffer (Takara, Shiga, Japan), 1 U of Ex *Taq* DNA polymerase, 4 nmol of dNTP mixture, 10 pmol of rHSP70_5RACE2 primer and GeneRacer 5’ primer (Table 1). The amplified products were diluted 10-fold with sterile distilled water and used as the template for the nested PCR with rHSP70_5RACE3 and GeneRacer 5’ nested primers (supplementary table S1). The PCR conditions were same as the first round PCR except for the template and primers. The amplified products were subcloned into pGEM-T vector (Promega, Madison, WI) and sequenced as reported previously (Yoon et al. 2008).

3’ RACE for rotifer HSP70 gene was carried out as follows. An aliquot of first strand cDNA was synthesized from rotifers in an exponential phase using an oligo-dT adapter primer (supplementary table S1). PCR was performed with the first strand cDNA as a template and with rHSP70_3RACE1 and AUAP primers. PCR was carried out with 30 cycles of 94°;C for 30 s, 60°;C for 30 s, and 72°;C for 2 min. The final extension step was performed at 72°;C for 5 min. The 20 µl of reaction mixture contained 5 pmol of forward and reverse primers, approximately 1 µg of first strand cDNA, 4 nmol of dNTP mixture, 2 µL of 10 × PCR buffer [100 mM Tris-HCl, pH8.3, 500 mM KCl, 15 mM MgCl2, 0.01% (w/v) gelatin] and 0.2 U of Ex *Taq* DNA polymerase (Takara). Nested PCR was performed under the same conditions using the PCR product diluted 100-fold and the rHSP70_3RACE2 primer. The amplified products were subcloned into pGEM-T vecter (Promega) and sequenced as described above. The nucleotide sequence of the open reading frame was confirmed by a single PCR using rHSP70_full_f1 and rHSP70_full_r1 primers (supplementary table S1). Conditions of the single PCR were same as described in 5’ RACE.

### Semi-Quantitative RT-PCR

RNA extraction and 1st strand cDNA synthesis were performed as described above. Primers rHSP70_gapF and rHSP70_gapR were designed to amplify the region encoding the GGMP repeat of *B. plicatilis* HSP70cB1i and HSP70cB2i genes (supplementary table S1). The b-actin gene was used as the internal control with specific primers rActinF and rActinR reported previously (Kaneko et al. 2005). PCR was carried out at 94°;C for 3 min followed by 20 – 40 cycles of 94°;C for 30 s, 58°;C for 30 s and 72°;C for 30 s. A 20-µL reaction mixture contained approximately 1 µg of 1st strand cDNA, 10 pmol of rHSP70_gapF and rHSP70_gapR, 2 µL of Ampli *Taq* buffer (Applied Biosystems, Foster City, CA), 4 nmol of dNTP mixture, and 0.2 U of Ampli *Taq* DNA polymerase (Applied Biosystems). The PCR products were separated with an 8% acrylamide gel and visualized by ethidium bromide staining.

### Quantitative Real-Time PCR

Rotifers were collected 0, 2, 4, 6, and 8 h after the heat treatment, and stored in ISOGEN (Takara) at -80°;C until use (three replicates for each time point, one replicate contained eight individuals). Rotifers collected before the heat treatment were used as the control. Quantitative real-time PCR was performed as reported previously (Kaneko et al. 2011). Primers for the b-actin gene were described previously (Kaneko et al. 2005); primers for HSP70 genes were designed using the Primer Express software ver. 2 (Applied Biosystems) (supplementary table S1). R version 3.5.1. and the multcomp package (Hothorn et al. 2008) was used for one-way analysis of variance (ANOVA) followed by the Dunnett’s test on the Macintosh platform.

### *In situ* Hybridization

Rotifers were fixed in PBS containing 4% paraformaldehyde 4 h after the heat treatment. *In situ* hybridization was performed as reported previously (Kaneko et al. 2011), using DIG-labeled RNA probes synthesized as follows. Primers HSP70_insituF and HSP70_insituR were designed to amplify the DNA fragment of 2026 - 2103 nt of *B. plicatilis* HSP70cB1i gene (supplementary table S1). PCR was performed in a 20 µL reaction mixture containing approximately 1 µg of the genomic DNA of *B. plicatilis,* 10 pmol each of the HSP70_insituF and HSP70_insituR primers, 2 µL of 10 × PCR buffer, 4 nmol dNTP mixture, and 0.2 U of Ex *Taq* DNA polymerase (Takara). The amplification was carried out with the initial denaturation at 94°;C for 3 min and 40 cycles of 94°;C for 30 s, 60°;C for 30 s, and 72°;C for 1.5 min, followed by the final extension step at 72°;C for 5 min. The amplified products were subcloned into pGEM-T easy vector (Promega). DIG-labeled RNA probes were synthesized with a DIG RNA labeling kit (Roche) according to the manufacturer’s instructions.

## References

Armougom F, Moretti S, Poirot O, Audic S, Dumas P, Schaeli B, Keduas V, Notredame C. 2006. Expresso: automatic incorporation of structural information in multiple sequence alignments using 3D-Coffee. Nucleic Acids Res 34:W604–W608.

Baird NA, Turnbull DW, Johnson EA. 2006. Induction of the heat shock pathway during hypoxia requires regulation of heat shock factor by hypoxia-inducible factor-1. J Biol Chem 281:38675–38681.

Boorstein WR, Ziegelhoffer T, Craig EA. 1994. Molecular evolution of the HSP70 multigene family. J Mol Evol 38:1–17.

Ceyhun SB, Sentürk M, Ekinci D, Erdoğan O, Ciltaş A, Kocaman EM. 2010. Deltamethrin attenuates antioxidant defense system and induces the expression of heat shock protein 70 in rainbow trout. *Comp Biochem Physiol*, C Toxicol Pharmacol 152:215–223.

Cottin D, Ravaux J, Leger N, Halary S, Toullec JY, Sarradin PM, Gaill F, Shillito B. 2008. Thermal biology of the deep-sea vent annelied *Paralvinella grasslei*: in vivo studies. J Exp Biol 211:2196–2204.

Daugaard M, Rohde M, Jäättelä M. 2007. The heat shock protein 70 family: Highly homologous proteins with overlapping and distinct functions. FEBS Lett 581:3702–3710.

De Nadal E, Ammerer G, Posas F. 2011. Controlling gene expression in response to stress. Nat Rev Genet 12:833–845.

Demand J, Lüders J, Höhfeld J. 1998. The carboxy-terminal domain of Hsc70 provides binding sites for a distinct set of chaperone cofactors. Mol Cell Biol 18:2023–2028.

Florin L, Becker KA, Sapp C, Lambert C, Sirma H, Müller M, Streeck RE, Sapp M. 2004. Nuclear translocation of papillomavirus minor capsid protein L2 requires Hsc70. J Virol 78:5546–5553.

Fröbius AC, Funch P. 2017. Rotiferan Hox genes give new insights into the evolution of metazoan bodyplans. Nat Commun 8:9.

Garbuz D. 2017. Regulation of heat shock gene expression in response to stress. Mol Biol 51:352–367.

Garbuz DG, Yushenova IA, Zatsepina OG, Przhiboro AA, Bettencourt BR, Evgen’ev MB. 2011. Organization and evolution of hsp70 clusters strikingly differ in two species of Stratiomyidae (Diptera) inhabiting thermally contrasting environments. BMC Evol Biol 11:74.

Hartl FU, Bracher A, Hayer-Hartl M. 2011. Molecular chaperones in protein folding and proteostasis. Nature 475:324–332.

Hess K, Oliverio R, Nguyen P, Le D, Ellis J, Kdeiss B, Ord S, Chalkia D, Nikolaidis N. 2018. Concurrent action of purifying selection and gene conversion results in extreme conservation of the major stress-inducible Hsp70 genes in mammals. Sci Rep 8:1–16.

Jayasena S, Chandrasekharan N, Karunanayake Physiological consequences of the supralittoral fringe: microhabitat temperature profiles and stress protein levels in the tropical periwinkle Cenchritis muricatus (Linneaus, 1758). Hydrobiologia 675:143–156.

EH. 1999. Molecular characterisation of a hsp70 gene from the filarial parasite *Setaria digitata*. Int J Parasitol 29:581–591.

Jedlicka P, Mortin MA, Wu C. 1997. Multiple functions of *Drosophila* heat shock transcription factor *in vivo*. EMBO J 16:2452–2462.

Judge ML, Botton ML, Hamilton MG. 2011. Physiological consequences of the supralittoral fringe: microhabitat temperature profiles and stress protein levels in the tropical periwinkle Cenchritis muricatus (Linneaus, 1758). Hydrobiologia 675:143-156.

Kaneko G, Yoshinaga T, Gribble KE, Mark Welch D, Ushio H. 2016. Measurement of survival time in *Brachionus* rotifers: synchronization of maternal conditions. J Vis Exp 113:e54126.

Kourtidis A, Drosopoulou E, Nikolaidis N, Hatzi VI, Chintiroglou CC, Scouras ZG. 2006. Identification of several cytoplasmic HSP70 genes from the Mediterranean mussel (*Mytilus galloprovincialis*) and their long-term evolution in Mollusca and Metazoa. J Mol Evol 62:446–459.

Le SQ, Gascuel O. 2008. An improved general amino acid replacement matrix. Mol Biol Evol 25:1307–1320.

Liu L, Cheng T-y, Yang Y. 2017. Cloning and expression pattern of a heat shock cognate protein 70 gene in ticks (*Haemaphysalis flava*). Parasitol Res 116:1695–1703.

Lo W-Y, Liu K-F, Liao I-C, Song Y-L. 2004. Cloning and molecular characterization of heat shock cognate 70 from tiger shrimp (*Penaeus monodon*). Cell Stress Chaperones 9:332.

Luan W, Li F, Zhang J, Wen R, Li Y, Xiang J. 2010. Identification of a novel inducible cytosolic Hsp70 gene in Chinese shrimp *Fenneropenaeus chinensis* and comparison of its expression with the cognate Hsc70 under different stresses. Cell Stress Chaperones 15:83–93.

Marchler G, Wu C. 2001. Modulation of *Drosophila* heat shock transcription factor activity by the molecular chaperone DROJ1. EMBO J 20:499–509.

Miernyk JA. 1997. The 70 kDa stress-related proteins as molecular chaperones. Trends Plant Sci 2:180–187.

Miller MA, Pfeiffer W, Schwartz T. 2010. Creating the CIPRES Science Gateway for inference of large phylogenetic trees. Proceedings of the Gateway Computing Environments Workshop (GCE); New Orleans, LA. p. 1–8.

Nikolaidis N, Nei M. 2004. Concerted and nonconcerted evolution of the Hsp70 gene superfamily in two sibling species of nematodes. Mol Biol Evol 21:498–505.

Piano A, Asirelli C, Caselli F, Fabbri E. 2002. Hsp70 expression in thermally stressed *Ostrea edulis*, a commercially important oyster in Europe. Cell Stress Chaperones 7:250.

Pollock DD, Larkin JC. 2004. Estimating the degree of saturation in mutant screens. Genetics 168:489–502.

Sanders BM. 1993. Stress proteins in aquatic organisms - an environmental perspective. Crit Rev Toxicol 23:49–75.

Sievers F, Wilm A, Dineen D, Gibson TJ, Karplus K, Li W, Lopez R, McWilliam H, Remmert M, Söding J. 2011. Fast, scalable generation of high-quality protein multiple sequence alignments using Clustal Omega. Mol Syst Biol 7:539.

Simoncelli F, Morosi L, Di Rosa I, Pascolini R, Fagotti A. 2010. Molecular characterization and expression of a heat-shock cognate 70 (Hsc70) and a heat-shock protein 70 (Hsp70) cDNAs in Rana (Pelophylax) lessonae embryos. *Comp Biochem Physiol*, A Mol Integr Physiol 156:552–560.

Sørensen JG, Kristensen TN, Loeschcke V. 2003. The evolutionary and ecological role of heat shock proteins. Ecol Lett 6:1025–1037.

Sorger PK, Pelham HR. 1988. Yeast heat shock factor is an essential DNA-binding protein that exhibits temperature-dependent phosphorylation. Cell 54:855–864.

Struck TH, Wey-Fabrizius AR, Golombek A, Hering L, Weigert A, Bleidorn C, Klebow S, Iakovenko N, Hausdorf B, Petersen M. 2014. Platyzoan paraphyly based on phylogenomic data supports a noncoelomate ancestry of Spiralia. Mol Biol Evol 31:1833–1849.

Touchon M, Hoede C, Tenaillon O, Barbe V, Baeriswyl S, Bidet P, Bingen E, Bonacorsi S, Bouchier C, Bouvet O. 2009. Organised genome dynamics in the *Escherichia coli* species results in highly diverse adaptive paths. PLoS genet 5:e1000344.

Wallace IM, O’sullivan O, Higgins DG, Notredame C. 2006. M-Coffee: combining multiple sequence alignment methods with T-Coffee. Nucleic Acids Res 34:1692–1699.

Yabu T, Imamura S, Mohammed MS, Touhata K, Minami T, Terayama M, Yamashita M. 2011. Differential gene expression of HSC70/HSP70 in yellowtail cells in response to chaperone- mediated autophagy. FEBS J 278:673–685.

Yoshinaga T, Minegishi Y, Rumengan IFM, Kaneko G, Furukawa S, Yanagawa Y, Tsukamoto K, Watabe S. 2004. Molecular phylogeny of the rotifers with two Indonesian *Brachionus* lineages. Coast Mar Sci 29:45–56.

Zheng G, Dong S, Hou Y, Yang K, Yu X. 2012. Molecular characteristics of HSC70 gene and its expression in the golden apple snails, *Pomacea canaliculata* (Mollusca: Gastropoda). Aquaculture 358–359:41-49.

Zuiderweg ER, Hightower LE, Gestwicki JE. 2017. The remarkable multivalency of the Hsp70 chaperones. Cell Stress Chaperones 22:173–189.

## References

Hothorn T, Bretz F, Westfall P. 2008. Simultaneous inference in general parametric models. Biom J 50:346–363.

Kaneko G, Kinoshita S, Yoshinaga T, Tsukamoto K, Watabe S. 2002. Changes in expression patterns of stress protein genes during population growth of the rotifer *Brachionus plicatilis*. Fish Sci 68:1317–1323.

Kaneko G, Yoshinaga T, Yanagawa Y, Kinoshita S, Tsukamoto K, Watabe S. 2005. Molecular characterization of Mn-superoxide dismutase and gene expression studies in dietary restricted *Brachionus plicatilis* rotifers. Hydrobiologia 546:117–123.

Kaneko G, Yoshinaga T, Yanagawa Y, Ozaki Y, Tsukamoto K, Watabe S. 2011. Calorie restriction-induced maternal longevity is transmitted to their daughters in a rotifer. Funct Eco/25:209-216.

Yoon SH, Itoh Y, Kaneko G, Nakaniwa M, Ohta M, Watabe S. 2008. Molecular characterization of Japanese sillago vitellogenin and changes in its expression levels on exposure to 17 beta-estradiol and 4-tert­ octylphenol. Mar Biotechno: 10 19–30.

